# Integrated Computational and Spectroscopic Analyses to unravel the Role of Cu(II) Coordination in Modulating Doxorubicin-DNA Binding

**DOI:** 10.64898/2026.09.11.750536

**Authors:** Dipanshu Ranjan Pattanayak, Aditya Dileep Kurdekar, Chelli Sai Manohar, R Sarojini, R Dharmaraj

**Affiliations:** Department of Physics, Sri Sathya Sai Institute of Higher Learning (SSSIHL), Prasanthi Nilayam, Sri Sathya Sai District, Andhra Pradesh, 515134, India; Department of Chemistry, SRM University, 5th Mile, Tadong, Gangtok, 737102, India; Independant Researcher, Coimbatore, Tamil Nadu, 641046, India

**Keywords:** Doxorubicin–DNA intercalation, Cu(II) mediation, Spectroscopy

## Abstract

Doxorubicin (DOX) is a clinically important anthracycline whose anticancer activity is closely associated with DNA binding. However, the structural basis by which metal coordination modulates DOX-DNA association remains poorly understood. Here, an integrated computational and spectroscopic approach was employed to elucidate the role of Cu(II) coordination in modulating the structure of DOX and its DNA binding. DFT, molecular docking, and molecular dynamics simulations established a stable intercalative association of DOX with DNA. At the same time, DFT-based characterization of Cu(II)-DOX revealed coordination-induced structural and electronic changes consistent with reduced conformational freedom and increased rigidity. This led to the hypothesis that Cu(II) coordination may favor an intercalation-compatible configuration of DOX. UV–Vis and competitive fluorescence studies experimentally supported Cu(II)-dependent modulation of DOX– DNA association, with UV–Vis analysis indicating a substantial tenfold increase in apparent DNA-binding affinity and fluorescence measurements confirming pronounced perturbation of DNA-associated ethidium bromide. Collectively, the findings establish a coherent relationship between Cu(II) coordination, DOX conformational modulation, and DNA recognition, highlighting metal coordination as a potential strategy for tuning the biomolecular interactions of anthracycline therapeutics.

## 1. Introduction

Deoxyribonucleic acid (DNA) is a fundamental biological macromolecule and a key target for therapeutic molecules, enabling molecular recognition through electrostatic interactions, hydrogen bonding, groove binding, and intercalation [1]. Doxorubicin (DOX) is a widely used anthracycline anticancer drug for treating various solid tumors and hematological malignancies (Fig. 1), exerting its anticancer activity primarily through DNA damage and inhibition of cancer cell proliferation[2,3]. Its biological activity involves multiple molecular mechanisms, including DNA intercalation and interaction with topoisomerase-II. The planar anthracycline chromophore of DOX intercalates between adjacent DNA base pairs. At the same time, the daunosamine sugar moiety is positioned within the minor groove and contributes to interactions with neighbouring DNA residues [2,4]. In addition to direct DNA intercalation, DOX acts as a topoisomerase-II poison by stabilizing the topoisomerase II–DNA cleavage complex and preventing efficient religation of cleaved DNA [3,5]. These mechanisms contribute to DNA damage and ultimately to cancer-cell death. Understanding the molecular determinants governing DOX–DNA interaction is crucial for elucidating its mechanism of action and for developing strategies to modify its physicochemical and biological properties. Despite its clinical importance, the therapeutic use of DOX is associated with limitations such as dose-dependent toxicity and the development of resistance [6,7]. These limitations have stimulated interest in approaches that modify the properties and biomolecular interactions of established anticancer agents.

**Fig. 1.**
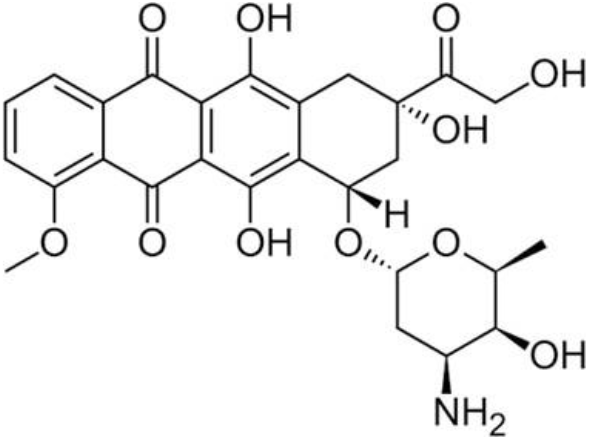
Two-dimensional chemical structure of DOX, showing its characteristic anthracycline framework.

Metal-based compounds have gained considerable importance in cancer chemotherapy because metal centers can provide distinctive coordination geometries, electronic properties, redox behavior, and modes of interaction with biomolecular targets [8,9]. The clinical success of platinum-based drugs such as cisplatin, carboplatin, and oxaliplatin, together with the therapeutic application of arsenic trioxide (As₂O₃), demonstrates the potential of metal-containing compounds in cancer treatment[8–10]. These examples have stimulated interest in other biologically relevant metal ions for the development of new therapeutic systems and the modification of existing drug properties.

Among biologically relevant transition metals, Cu(II) is particularly attractive because of its essential biological role and versatile coordination chemistry. Copper participates in electron-transfer reactions, enzyme catalysis, redox regulation, and cellular signalling, while its flexible coordination behaviour enables interactions with a wide range of ligands and biomolecular targets[11–13]. Coordination of Cu(II) with an organic drug can modify its molecular geometry, charge distribution, electronic properties, lipophilicity, stability, and interactions with biological macromolecules[11,12]. However, the biological behaviour of Cu(II)-containing systems depends strongly on their coordination environment, stability, and interactions with the surrounding biological medium. Understanding the structural and molecular consequences of Cu(II) coordination to a bioactive drug is essential for evaluating its influence on biomolecular interactions [14].

In the case of DOX, coordination with Cu(II) provides a potentially important means of modifying its interaction with DNA. Complex formation between DOX and Cu(II) can alter the electronic distribution and molecular geometry of the drug, thereby influencing its ability to interact with DNA [15–17]. Structural changes in the anthracycline chromophore and restriction of its conformational flexibility affect the geometric complementarity required for intercalation. If Cu(II) coordination reduces the number of accessible conformational degrees of freedom, the resulting complex may be better predisposed to adopt and maintain an intercalative orientation within the DNA base-pair stack. Thus, Cu(II) coordination may also influence the conformational and structural characteristics that govern its recognition by DNA.

Although the interactions of DOX with DNA and the therapeutic potential of metal-based anticancer agents have been extensively studied, the molecular role of Cu(II) coordination in modulating the structure, electronic properties, and atomistic insights into DOX’s DNA-intercalative behavior remain insufficiently understood. In particular, how Cu(II) coordination alters DOX and its DNA-binding behavior remains unclear, warranting further investigation into the structural and electronic changes underlying the DOX-Cu(II)-DNA interaction [15,17–22].

In the present study, the DOX-DNA system and the role of Cu(II) coordination are investigated through a combination of computational and experimental approaches. Docking and 100 ns molecular dynamics simulations demonstrate that DOX maintains a stable intercalative configuration within DNA, providing a molecular reference for evaluating the effects of Cu(II) coordination. DFT analyses further reveal distinct Cu(II)-induced structural and electronic changes in DOX, particularly in features relevant to its DNA-binding configuration, while UV–Vis absorption and fluorescence measurements demonstrate corresponding changes in its DNA-binding characteristics. Collectively, these findings indicate that Cu(II) coordination reorganizes the structural and electronic features of DOX, thereby modulating its interaction with DNA and potentially influencing its intercalation behavior. By linking metal-induced molecular changes in DOX to its altered DNA-binding properties, this study provides a mechanistic perspective on how Cu(II) coordination can regulate DOX’s DNA-recognition behavior.

## 2. Methods and Materials

### 2.1 Computational Details

All quantum-chemical calculations were performed using Gaussian 16 [23]. The geometry optimization and frequency calculations of DOX were initially performed at the Hartree–Fock (HF) level, followed by Density Functional Theory (DFT) calculations using the Becke three-parameter Lee–Yang–Parr (B3LYP) functional along with the 6-311+G(d,p) basis set [24]. Grimme’s D3 dispersion correction (empirical dispersion = gd3) was included to account for dispersion interactions [25]. Solvent effects were incorporated using the SMD implicit solvation model for water [26]. All structures were fully optimized without symmetry constraints, and subsequent frequency calculations confirmed that the optimized geometries correspond to true minima on the potential energy surface, as indicated by the absence of imaginary frequencies. The vibrational frequency calculations were further used to obtain the IR and Raman spectra. Time-dependent density functional theory (TD-DFT) calculations were performed on the optimized geometries at the same level of theory (B3LYP/6-311+G(d,p) with SMD water model and D3 dispersion correction). All TD-DFT calculations were performed as vertical excitations from the ground-state-optimized geometries, enabling estimation of excitation energies and simulation of UV–Vis absorption spectra. For the DOX–Cu complex, the large system size and high computational cost necessitated a bilayer ONIOM approach. The inner core region, comprising the Cu center and directly coordinating oxygen atoms, was treated at the DFT level using the B3LYP functional with the LANL2DZ basis set. In contrast, the remaining part of the system was treated using the semi-empirical PM6 method [27]. The geometry of the complex was optimized at this ONIOM level, followed by frequency calculations to confirm the stability of the optimized structure and to obtain the corresponding IR and Raman spectra. The Multiwfn 3.8 program [28], along with VMD 1.9.3 [29], was used to analyze and visualize the NCI analysis.

The 3D structure of the DNA hexamer containing an intercalation gap (PDB ID: 1Z3F) was obtained from the RCSB Protein Data Bank [30]. The structure was cleaned by removing solvent molecules, metal ions, and any co-crystallized ligands. The prepared DNA structure was then converted to PDBQT format using Meeko, in which polar hydrogens were added, and partial atomic charges were assigned using the Kollman united-atom model and the Gasteiger–Marsili method. The optimized structure of DOX was prepared separately and converted into PDBQT format using the same procedure to ensure consistency in atom typing and charge assignment. Molecular docking studies were performed using AutoDock Vina [31]. The binding site was defined based on the position of the pre-existing intercalated ligand in the crystal structure. Accordingly, a grid box was centered at X = 0.893, Y = 17.494, and Z = 46.285, with dimensions of 20×20×20 Å, covering the intercalation region. A semi-flexible docking approach was employed, where DOX was treated as a flexible ligand with rotatable bonds, while the DNA receptor was kept rigid. The docking poses were ranked by binding affinity and analyzed with PyMOL to examine the binding mode and key interactions [32].

From molecular docking, the best binding pose was selected as the initial configuration for molecular dynamics (MD) simulations. All MD simulations were performed using GROMACS 2026.0 [33]. The DNA was described using the AMBER99SB-ILDN force field [34], while the topology and force-field parameters for DOX were generated using ACPYPE [35], which provides AMBER-compatible parameters. The DNA-DOX complex was placed in a cubic simulation box with a minimum distance of 1.0 nm from the box edges and solvated using the SPC explicit water model. Sodium (Na⁺) counterions were added to neutralize the system’s net charge while maintaining periodic boundary conditions. The solvated system was first subjected to energy minimization using the steepest-descent algorithm until convergence, to remove unfavorable steric contacts and inappropriate geometric factors. Subsequently, the system was equilibrated for 100 ps under the NVT ensemble, followed by 500 ps under the NPT ensemble. Temperature was maintained at 300 K using the velocity-rescaling (V-rescale) thermostat [36]. In contrast, pressure was maintained at 1 bar using the Parrinello–Rahman barostat [37] owing to their reliable temperature and pressure control and appropriate ensemble sampling. Following equilibration, a 100 ns production MD simulation was performed with a 2-fs integration time step under periodic boundary conditions. Long-range electrostatic interactions were treated using the Particle Mesh Ewald (PME) method [38], which efficiently accounts for long-range Coulombic interactions under periodic boundary conditions with a 1.0 nm cutoff. In comparison, Van der Waals interactions were treated using a 1.0 nm cutoff throughout the simulation. All covalent bonds involving hydrogen atoms were constrained using the LINCS algorithm [39], thereby enabling a 2 fs time step throughout the simulation. Trajectory analyses were performed using the built-in analysis modules of GROMACS. The structural stability and dynamic behavior of the DNA–DOX complex were evaluated through the calculation of the root-mean-square deviation (RMSD), root-mean-square fluctuation (RMSF), radius of gyration (R_g_), minimum intermolecular distance, solvent-accessible surface area (SASA), and the number of intermolecular contacts over the 100 ns simulation.

### 2.2 Experimental Details

Herring sperm deoxyribonucleic acid (DNA) sodium salt and ethidium bromide (EB) of purity >98% were purchased from HiMedia Laboratories Pvt. Ltd., India. Doxorubicin hydrochloride (DOX) was obtained from Dabur Pharma Ltd., India. Copper (II) chloride (CuCl₂) was of analytical grade. All reagents were used as received without further purification. All solutions used for spectrophotometric measurements were prepared at room temperature in a 10 mM phosphate buffer (pH 7.0). A stock solution of herring sperm DNA was prepared by dissolving the required amount of DNA in phosphate buffer at room temperature, followed by occasional stirring for 24 h to ensure complete dissolution and homogeneity. The purity of the DNA preparation was assessed spectrophotometrically by determining the absorbance ratio at 260 and 280 nm (A_260_ /A_280_). The observed ratios were in the range of 1.8-1.9, indicating that the DNA was sufficiently free of protein contamination; therefore, no further deproteinization was performed [20,40]. The concentration of the DNA stock solution was determined from its absorbance at 260 nm using a molar extinction coefficient of ε_260_ = 6600 M⁻ ¹ cm⁻ ¹ per base [20].

A 0.5 mM stock solution of EB and a 2 mM stock solution of DOX were prepared by directly dissolving the respective compounds in a 10 mM phosphate buffer (pH 7.0). The stock solutions were stored at 4 °C and used within two days after preparation. All working solutions were prepared by appropriate dilution of the stock solution in 10 mM phosphate buffer (pH 7.0) at room temperature.

#### 2.2.1 Spectrophotometric Studies

UV–Vis absorption measurements were carried out using a Jasco V-530 UV–Vis spectrophotometer with a slit width of 2 nm, scan rate of 100 nm min⁻¹, and data pitch of 1 nm. Quartz cuvettes with a 1 cm path length were used throughout the measurements. The absorption spectra of DOX and EB were recorded in the wavelength ranges of 450–550 nm and 200–500 nm, respectively. Prior to recording the spectra, baseline correction was performed using a 10 mM phosphate buffer (pH 7.0). In all absorption experiments, the ligand, ligand–Cu(II), ligand–DNA, and ligand–Cu(II)–DNA systems were taken in the sample cuvette against phosphate buffer in the reference cuvette.

For the interaction studies between DOX and DNA, DOX was maintained at a fixed concentration of 80 μM and titrated with increasing DNA concentrations using a micro-injector. After each DNA addition, the solution was thoroughly mixed and allowed to equilibrate for 2 minutes before recording the absorption spectrum. DNA titration was continued until no further change in absorbance was observed, indicating saturation of the interaction. The total accumulated volume in the cuvette was maintained below 3 mL. The concentration of free DOX was selected such that its absorbance remained below 1 to comply with the Beer–Lambert law [41,42].

The interaction of DOX with Cu(II) was investigated by maintaining the DOX concentration at 80 μM and gradually increasing the Cu(II) concentration until saturation. To investigate the influence of Cu(II) on DOX–DNA binding, DOX–Cu(II) mixtures were prepared at molar ratios of 1:0.5, followed by titration with increasing concentrations of DNA. The resulting absorption spectra were recorded after equilibration until saturation was attained. All measurements were performed at room temperature.

The absorption titration data for the DOX–DNA interaction were fitted to the double-reciprocal equation 2.1 [43,44],

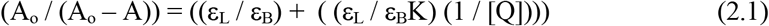

where A_o_ and A are the absorbances of DOX in the absence and presence of DNA, respectively; εL and ε_B_ are the molar extinction coefficients of free and bound DOX, respectively; [Q] is the DNA concentration; and K is the binding constant. The primary data were used to plot a linear double-reciprocal plot of (1/ (A_o_ – A)) versus (1/[Q]). Linear regression best fit analysis was used to estimate the binding constant, K, from the ratio of the intercept to slope, where the regression coefficient, r^2^, was 0.9 < r^2^ < 1.0.

The Gibbs free energy change, ΔG, associated with the ligand–DNA interaction was calculated from the binding constant obtained from the absorption titration using the thermodynamic relationship given in equation 2.2.

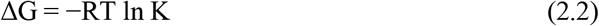

where R is the universal gas constant (8.314 J mol^-1^ K^-1^), T is the absolute temperature in kelvin, and K is the binding constant obtained from the double-reciprocal analysis. The calculated ΔG values were expressed in kJ mol^-1^.

#### 2.2.2 Spectrofluorimetric Studies

Fluorescence emission measurements were performed using a Jasco FP-750 spectrofluorometer with excitation and emission slit widths of 5 nm, a scan rate of 250 nm min⁻¹, and a data pitch of 1 nm. The instrument’s response and sensitivity were set to medium. High-quality glass cuvettes with a 1 cm path length were used for all fluorescence measurements. Appropriate buffer blanks were subtracted by setting the instrument to autozero to correct for background fluorescence. For competitive binding studies, the fluorescent probe EB was mixed with DNA at an appropriate molar ratio to form the EB–DNA complex, which was subsequently titrated with DOX. [43,45]. The stock solutions of EB and DNA were diluted with 10 mM phosphate buffer (pH 7.0) to concentrations of 15 and 60 μM, respectively. EB was excited at its absorption maximum of 480 nm, and the emission spectra were recorded from 550 to 650 nm.

To investigate the influence of Cu(II) on the competitive binding of EB and DOX with DNA, Cu(II) was added to the EB–DNA mixture at an EB:DNA(II) molar ratio of 1:4:0.4. The resulting EB–DNA–Cu(II) mixtures were subsequently titrated with increasing concentrations of DOX. After each addition, the solution was thoroughly mixed and allowed to equilibrate for 2 minutes before the fluorescence spectrum was recorded. All measurements were performed at room temperature.

The titration data corresponding to EB-DNA-ligand interactions are fitted to the Stern-Volmer (SV) equation 2.3 [20,46], to obtain the quenching rate constant, K_q_, for the fluorescence quenching studies.

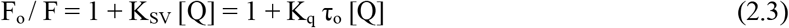

where, F_o_ and F are fluorescence intensities of the molecular fluorophore (EB-DNA or EB-DNA-metal ions) before and after the addition of the quencher (DOX).

[Q] is the concentration of the quencher (DOX)

K_sv_ is the Stern-Volmer quenching constant, and K_q_ is the quenching rate constant. τ_o_ is the fluorescence lifetime without quencher (DOX)

The fluorescence quenching titration data of EB-DNA-ligand are fitted to equation 2.4 [47] to calculate the binding parameters of EB with DNA on increasing the concentration of the quencher either in the absence or presence of metal ions.

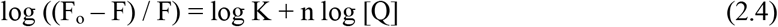

where,

F_o_ and F are fluorescence intensities of the molecular fluorophore (EB-DNA or EB-DNA-metal ions) before and after the addition of the quencher (DOX)

[Q] is the concentration of the quencher (DOX)

’K’ is the binding constant, and ’n’ is the binding stoichiometry or number of binding sites of DNA available for the molecular fluorophore.

The linear SV plot of F_o_ / F versus [Q] and the linear plot of log ((F_o_ ∼ F) / F) versus log [Q] were used along with the linear regression best fit analysis to estimate the value of K_q_, K, and n, where the regression coefficient value, r^2^, was 0.9 < r^2^ < 1.0.

For systems showing deviation from linear SV behaviour, the modified SV equation (Lehrer equation) given in equation 2.5 was used to determine the fraction of fluorophore accessible to the quencher and the binding constant for the accessible fraction [48,49].

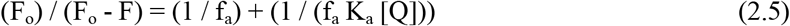

where F_o_ and F are the fluorescence intensities of the fluorophore in the absence and presence of the quencher, respectively; f_a_ is the fraction of fluorophore accessible to the quencher; K_a_ is the binding constant of the accessible fraction; and [Q] is the concentration of the quencher, DOX. A plot of (F_o_ / (F_o_-F)) versus (1/[Q]) was subjected to linear regression analysis. The intercept and slope of the linear plot were used to determine f_a_ and K_a_, respectively.

The fluorescence titration data were analysed using a nonlinear Hill binding equation [50] to evaluate the binding affinity and cooperativity of the ligand-DNA interaction, as described in equation 2.6,

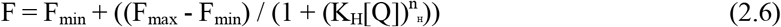

where F is the fluorescence intensity at a given DOX concentration, F_max_ and F_min_ represent the fluorescence intensities in the absence and at saturation of the ligand, respectively; K_H_ is the apparent binding constant; [Q] is the concentration of the ligand, and n_H_ is the Hill coefficient representing the apparent cooperativity of the interaction. The parameters K_H_ and n_H_ were obtained by nonlinear regression of the experimental fluorescence data. Values of (n_H_ > 1) and (n_H_ < 1) indicate positive and negative apparent cooperativity, respectively.

## 3. Results and Discussion

### 3.1 DFT Studies on DOX

The DFT-optimized structure of DOX (Fig. 2) revealed that the anthracycline chromophore adopts an approximately planar conformation, a structural feature that facilitates DNA intercalation through favorable π–π stacking interactions with base pairs. The optimized structure was subsequently employed for TD-DFT calculations, which predicted three principal electronic transitions at 448.75, 406.14, and 394.15 nm. Compared with the experimental absorption spectrum, a spectral deviation of approximately 31 nm is well within the range commonly encountered in TD-DFT calculations for extended aromatic and pharmaceutical systems. Such deviations can arise from several factors, including solvent effects, vibronic coupling, hydrogen-bonding interactions, and the inherent limitations of the exchange-correlation functional in accurately describing excited states, particularly its tendency to overestimate excitation energies. The most intense transition at 448.75 nm is predominantly assigned to a π→π* excitation within the anthracycline aromatic framework, involving promotion of electron density from occupied π orbitals to low-lying antibonding π* orbitals. The weaker transitions at 406.14 and 394.15 nm are attributed primarily to additional π→π* and n→π* excitations involving the conjugated aromatic and carbonyl-containing regions of DOX[51]. Overall, the calculated excitation profile reproduces the key features of the experimental absorption spectrum [52], and the optimized structure was used for subsequent electronic-structure and DNA-binding analyses.

**Fig. 2.**
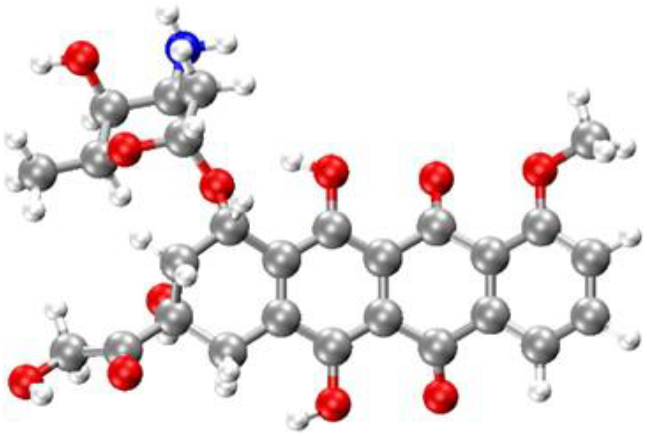
DFT-optimized molecular structure of doxorubicin (DOX) shown in a ball-and-stick representation. Carbon, oxygen, nitrogen, and hydrogen atoms are depicted in grey, red, blue, and white, respectively.

**Fig. 3.**
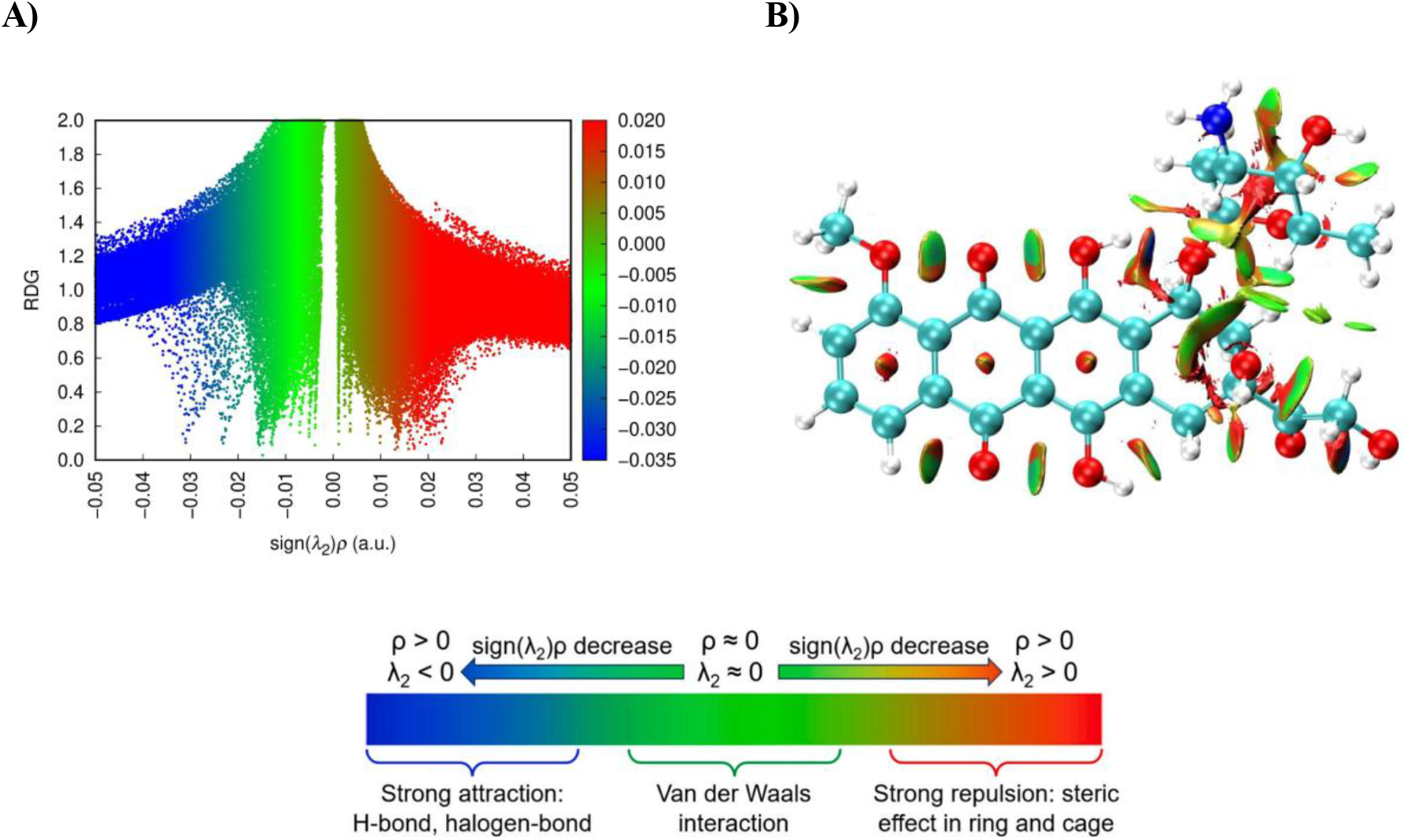
A) RDG scatter plot and B) NCI isosurface of the optimized DOX structure, illustrating the nature and spatial distribution of its noncovalent interactions. The interaction regions are characterized by sign(λ₂).ρ, where blue, green, and red features denote attractive, weak Van der Waals, and repulsive/steric interactions, respectively.

**Fig. 4.**
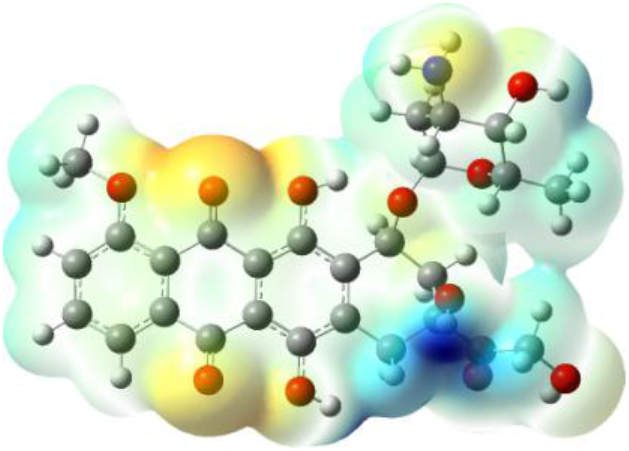
Molecular electrostatic potential (MESP) surface of the DFT-optimized DOX structure, illustrating the spatial distribution of electrostatic potential across the anthracycline and amino-sugar moieties. Electron-rich and electron-deficient regions are represented by negative and positive electrostatic potential, respectively, with intermediate colors corresponding to regions of moderate potential.

#### 3.1.1 NCI Analysis on DOX

The NCI analysis was performed to understand the various interactions present in DOX using the reduced density gradient (RDG) method [53]. This technique distinguishes among hydrogen-bonding interactions, Van der Waals forces, and steric effects across various parts of the molecule. The sign(λ_2_).ρ is the combination of electron density and the second Hessian eigenvalue sign. These interactions are categorized as attractive, repulsive, and intermediate, corresponding to sign(λ_2_).ρ < 0, sign(λ_2_).ρ > 0, and sign(λ_2_).ρ = 0, respectively. The three colors in the scatter plot and isosurfaces represent three different kinds of interactions: blue shades for favorable interactions, such as hydrogen bonding; yellow-green shades for Van der Waals forces; and red shades for repulsive interactions, such as steric effects. Using this method, a graphical representation of various zones of noncovalent interactions in real space was achieved.

The NCI analysis of DOX reveals a combination of weak dispersive, attractive, and steric interactions distributed over different structural regions of the molecule. The RDG scatter plot shows a pronounced concentration in the range −0.03 a.u. to 0.02 a.u. of the sign(λ₂).ρ value, which is consistent with the extensive green and red isosurfaces observed in the NCI isosurface. These green isosurfaces are distributed predominantly over the extended anthracycline framework, the common region shared by the sugar group, and the regions between adjacent functional groups, indicating that weak Van der Waals interactions contribute to the conformational stabilization of DOX. In particular, the broad green isosurfaces associated with the fused aromatic framework reflect the dispersive interactions present around the extended π-conjugated surface. In contrast, additional green regions are observed around the oxygen- and nitrogen-containing substituents and the amino-sugar portion of the molecule. The attractive component is represented by the negative sign(λ₂).ρ region of the scatter plot, with features extending to approximately −0.032 a.u. These interactions appear as blue-to-green regions on the NCI surface. They are primarily localized around the heteroatom-rich portions of DOX, particularly near the hydroxyl, carbonyl, and amino functionalities. These localized attractive regions indicate favorable noncovalent interactions involving polar functional groups and potential hydrogen bonding.

In contrast, the positive sign(λ₂).ρ region yields yellow-to-red isosurfaces and extends to approximately +0.025 a.u. in the RDG plot. The corresponding red regions are particularly evident toward the central portions of the fused aromatic rings and in geometrically congested regions around the substituted portions of the anthracycline and amino-sugar moieties. These regions arise due to steric repulsion within spatially confined regions of the molecule.

Thus, the NCI surface indicates that the relatively rigid fused-ring framework exhibits both dispersive interactions around the molecular surface and localized repulsive regions within the rings. At the same time, the peripheral functional groups contribute additional attractive interactions. Overall, the spatial distribution of these interactions provides a molecular-level description of the electronic and geometric factors governing the optimized DOX structure and establishes a suitable reference for comparison with the DOX-Cu²⁺ complex.

#### 3.1.2 MESP Analysis on DOX

The MESP surface of DOX reveals a distinctly heterogeneous distribution of electrostatic potential across the molecule. The electron-rich regions are predominantly localized around the oxygen-containing functional groups of the anthracycline framework, as evident from the yellow-to-orange regions surrounding the hydroxyl and carbonyl oxygen atoms. These regions indicate a relatively negative electrostatic potential representing the nucleophilic/electron-rich sites of DOX, particularly pronounced near the quinone/carbonyl-containing portion of the fused aromatic framework. This reflects the strong contribution of these heteroatoms to the overall electrostatic character of the molecule [54]. In contrast, a pronounced blue region is observed around the amino group of the amino-sugar moiety, indicating an electron-deficient/positive electrostatic potential in this part of the molecule.

The strong localization of positive potential around the nitrogen-containing region distinguishes the amino-sugar portion from the predominantly neutral-to-electron-rich regions of the anthracycline core. The spatial separation between the electron-rich oxygen-containing sites and the electron-deficient amino region demonstrates the heterogeneous electrostatic character and polarization of DOX. The extended anthracycline framework itself is dominated by green-to-cyan regions, corresponding to comparatively moderate electrostatic potential, with localized changes around its substituted functional groups. Thus, the MESP surface identifies the oxygen-containing functionalities and the amino group as the most electrostatically differentiated regions of DOX. This complementary distribution provides a molecular basis for favorable noncovalent interactions and is particularly relevant to the subsequent analysis of Cu²⁺ complexation, in which electron-rich oxygen sites can contribute to metal coordination, thereby modifying the electrostatic profile of DOX.

### 3.2 Docking Studies on DOX-DNA

Molecular docking of the DFT-optimized DOX structure with the DNA hexamer revealed a favorable binding interaction within the intercalation site. The best-ranked docking pose exhibited a binding affinity of −7.84 kcal/mol, indicating a stable ligand–DNA complex. The subsequent poses showed slightly higher energies, with small RMSD deviations, suggesting moderate conformational variability but a consistent binding region. The top-ranked pose clearly demonstrates that DOX adopts intercalative adjacent base pairs. As observed in Fig. 5, the ligand is positioned within the intercalation gap formed by the DNA duplex, stabilizing the complex through π–π stacking interactions with neighboring nucleobases, particularly guanine residues (DG). In addition to stacking interactions, several hydrogen bonds were observed between DOX and the DNA bases, with distances ranging from approximately 2.2 to 2.7 Å, indicating strong, directional hydrogen bonding. These interactions significantly stabilize the ligand within the DNA cavity. The involvement of both hydrogen bonding and π–π stacking highlights the dual nature of DOX binding. The relatively low RMSD values among the other poses suggest that the ligand consistently adopts a similar orientation within the intercalation site, reinforcing the reliability of the predicted binding mode. Overall, the docking results confirm that DOX preferentially binds to DNA via classical intercalation, driven by strong π–π stacking interactions and supported by hydrogen bonding with surrounding nucleobases. This binding mode is consistent with the known mechanism of action of DOX as a DNA-intercalating anticancer agent [55].

**Fig. 5.**
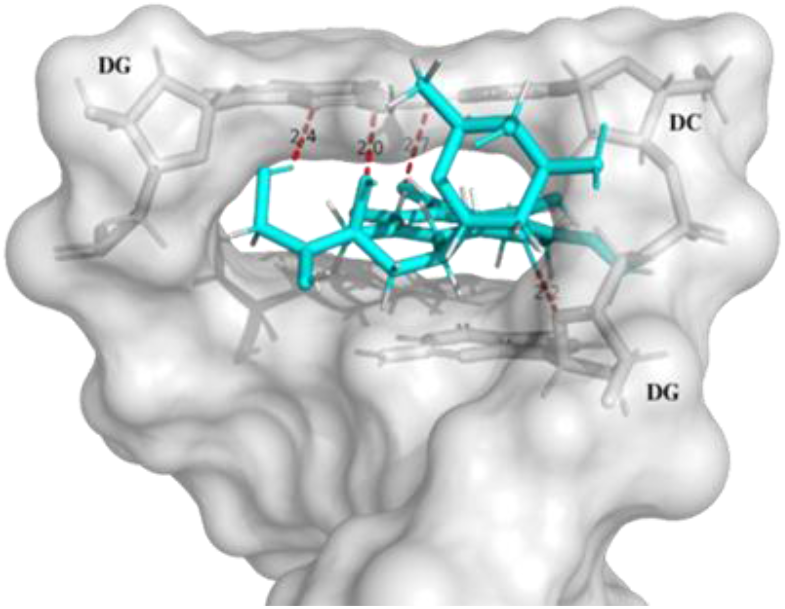
Molecular representation of DOX intercalated within the DNA hexamer. The DNA is shown as a grey Van der Waals surface, DOX as a cyan stick model, and hydrogen-bonding interactions as red dashed lines.

### 3.3 MD Studies on DOX-DNA

Molecular dynamics (MD) simulations have previously been employed to investigate the stability and intercalative behavior of DOX–DNA complexes[56]. Building on these established computational studies, the present 100 ns MD trajectory was analyzed using several complementary structural and interaction-based parameters, including RMSD, RMSF, radius of gyration (Rg), minimum intermolecular distance, SASA, and the number of intermolecular contacts to verify the structural stability and persistence of the docked DOX–DNA complex. The results obtained from each parameter are discussed below.

As shown in Fig. 6(A), the DNA backbone initially exhibited a rapid structural relaxation and then attained a stable conformation (∼ 0.20–0.24 nm). In comparison, DOX exhibited relatively larger fluctuations, primarily within the 0.20–0.30 nm range, with occasional transient increases during the simulation. These fluctuations are attributed to local conformational adjustments of the ligand within the DNA binding pocket rather than ligand dissociation. Importantly, the ligand RMSD remained stable over the entire 100 ns trajectory without any continuous drift, indicating that DOX remained firmly associated with the DNA duplex.

**Fig. 6.**
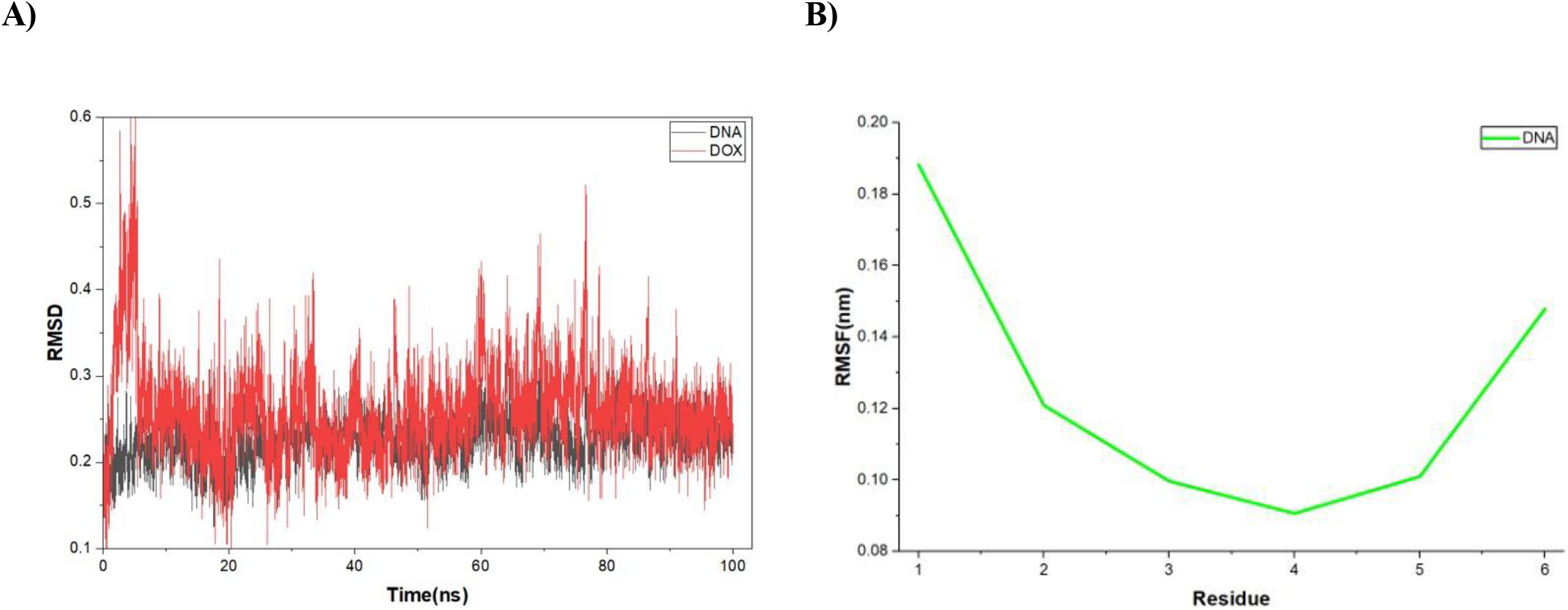
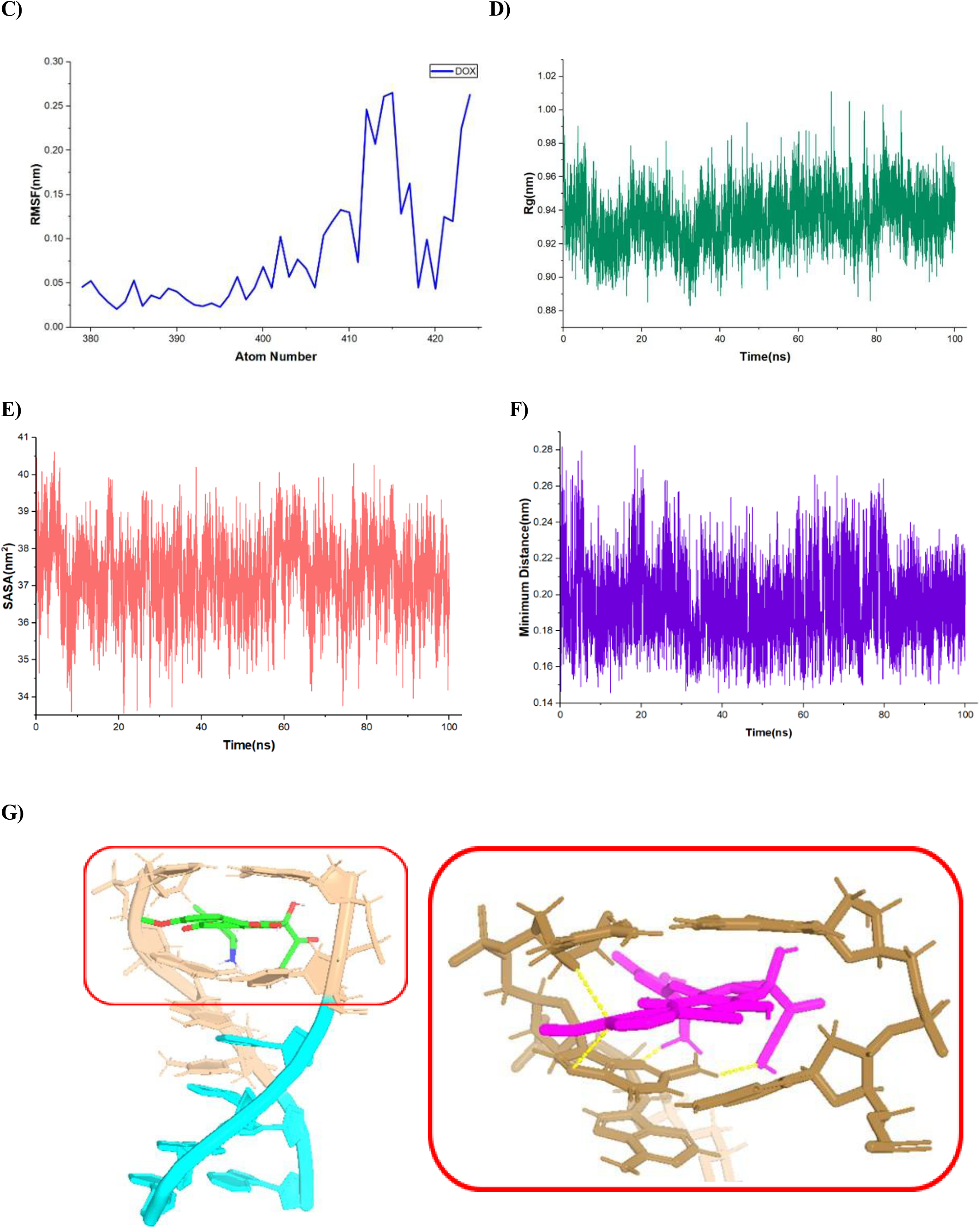
A) RMSD profiles of the DNA backbone (black) and DOX (red); B) residue-wise RMSF profile of the DNA duplex; C) atom-wise RMSF profile of DOX; D) radius of gyration (Rg) of the DNA–DOX complex; E) solvent-accessible surface area (SASA); F) minimum distance between DOX and DNA throughout the MD simulation; and G) representative DNA–DOX complex extracted from the MD trajectory, with enlarged views of the binding region showing the intercalated DOX and surrounding DNA bases.

Further, as observed in Fig. 6(B, C), the DNA exhibited relatively low residue-wise fluctuations (RMSF values: ∼ 0.09-0.19 nm). The terminal residues exhibited greater fluctuations than the central residues, as expected due to their greater conformational freedom. In contrast, the central region of the DNA remained comparatively rigid, indicating that the overall duplex structure was well preserved during the simulation. However, DOX revealed generally low atomic fluctuations for most atoms, with values < 0.10 nm. Slightly higher fluctuations were observed for atoms associated with the terminal functional groups and flexible side chains. These localized fluctuations reflect the intrinsic flexibility of the ligand rather than instability within the DNA binding site. Overall, the low flexibility of the DNA core together with the limited fluctuations of the ligand further supports the stable binding of DOX within the DNA duplex.

As shown in Fig. 6(D), the radius of gyration (R_g_) remained relatively stable throughout the simulation, fluctuating within a narrow range of approximately 0.90–0.97 nm. Minor fluctuations were observed during the trajectory, which can be attributed to the natural thermal motion of the DNA duplex in the solvated environment. To corroborate this, we also evaluated the minimum intermolecular distance, which remained relatively constant, fluctuating primarily between 0.18 and 0.22 nm, with only minor transient deviations due to thermal motion.

The solvent-accessible surface area (SASA) of the DNA–DOX complex was determined to study changes in solvent exposure and overall structural stability. As shown in Fig. 6(E), the SASA values fluctuated within a relatively narrow range of approximately 36–39 nm² during the entire simulation. These minor fluctuations are attributed to the inherent thermal motion of the solvated system. The absence of large variations suggests that the binding of DOX did not induce substantial structural expansion or contraction of the DNA duplex. Finally, an analysis of the 10001 trajectory frames identified eight distinct conformational clusters. Among these, the first cluster was overwhelmingly dominant, containing 9769 structures (≈97.7%) (SI Fig. S1). In addition, the average pairwise RMSD of 0.092 nm indicates minimal structural deviation among the sampled configurations. The predominance of a single conformational cluster indicates that the DNA–DOX complex rapidly converged on a stable structural state and remained in this energetically favorable conformation for nearly the entire simulation. The minor clusters most likely correspond to transient local structural fluctuations rather than significant conformational transitions. To visualize the binding mode after MD simulation, a representative structure from the equilibrated 100 ns trajectory was analyzed using PyMOL (Fig. 6(G)). Here, we observe that the DOX remained stably intercalated between adjacent DNA base pairs without significant displacement, while the DNA duplex retained its overall architecture. The close-up view shows stabilization through multiple noncovalent interactions with the surrounding nucleobases. This persistent intercalative mode is consistent with the stable RMSD, minimum distance, and intermolecular contact profiles, supporting sustained DOX–DNA binding throughout the simulation.

Overall, the MD analyses provide a consistent picture of the structural stability of the DNA–DOX complex over 100 ns (SI Video S1). Stable RMSD and RMSF profiles, preserved radius of gyration, minimum intermolecular distance, SASA, and persistent intermolecular contacts indicate that DOX remained stably bound within the DNA binding site without significant dissociation. These results support the stability of the predicted docking pose and the sustained DOX–DNA interactions under the simulated conditions.

### 3.4 DOX-Cu Complex

The DFT-optimized structure of the Cu–DOX complex revealed a distorted square-planar coordination geometry around the Cu(II) center, coordinated through the oxygen donor sites of DOX (Fig. 7), involving the carbonyl and hydroxyl functional groups of the anthracycline moiety. The optimized structure suggests stable Cu(II) chelation, resulting in a localized rearrangement of these donor groups around the metal center, while the extended anthracycline framework is largely retained. Thus, Cu(II) coordination primarily modifies the oxygen-rich coordination region and its local electronic environment [57].

**Fig. 7.**
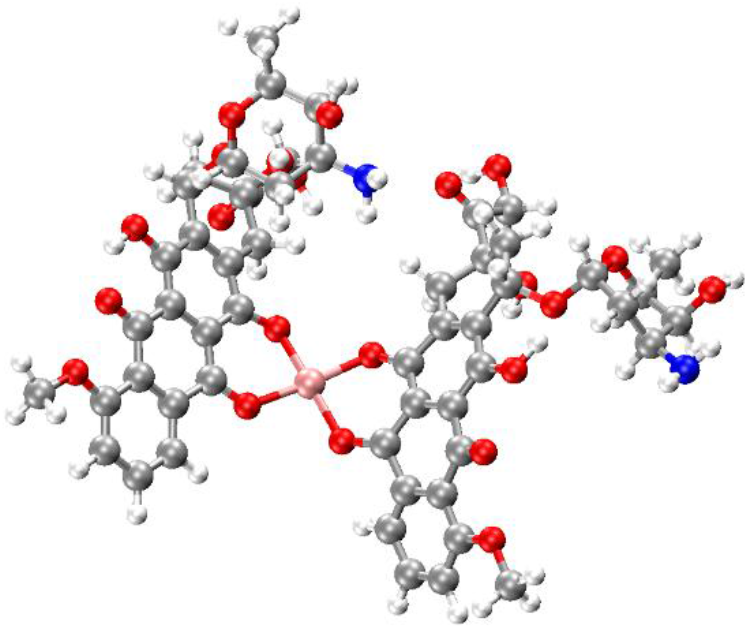
DFT optimized DOX-Cu complex shown in a ball-and-stick representation. Carbon, oxygen, nitrogen, copper, and hydrogen atoms are depicted in grey, red, blue, pink, and white, respectively.

To elucidate the changes in electronic structure and intermolecular interactions upon Cu(II) coordination, a comparative analysis of the NCI, MESP, and IR/Raman spectra of free DOX and the Cu–DOX complex is presented below.

#### 3.4.1 Comparative NCI Analysis

Compared with free DOX, the Cu–DOX complex exhibits additional localized NCI features around the Cu(II) coordination center and the surrounding oxygen donor atoms (Fig. 8). The RDG scatter plot shows attractive contributions extending to approximately sign(λ₂).ρ = −0.035 a.u., with the corresponding blue features appearing within the NCI isosurface, particularly along the regions connecting the Cu(II) center with the coordinated oxygen atoms. These features are consistent with the favorable attractive interactions associated with the Cu²⁺– O coordination environment involving the carbonyl and hydroxyl donor sites. The extensive green-to-yellow regions distributed around the anthracycline framework and between its oxygen-containing functional groups correspond to weak Van der Waals interactions, similar to those observed for free DOX. In addition, red features associated with positive sign(λ₂).ρ values, reaching approximately +0.020 a.u., are observed around the Cu coordination region and within sterically congested portions of the molecular framework, indicating localized repulsive interactions.

**Fig. 8.**
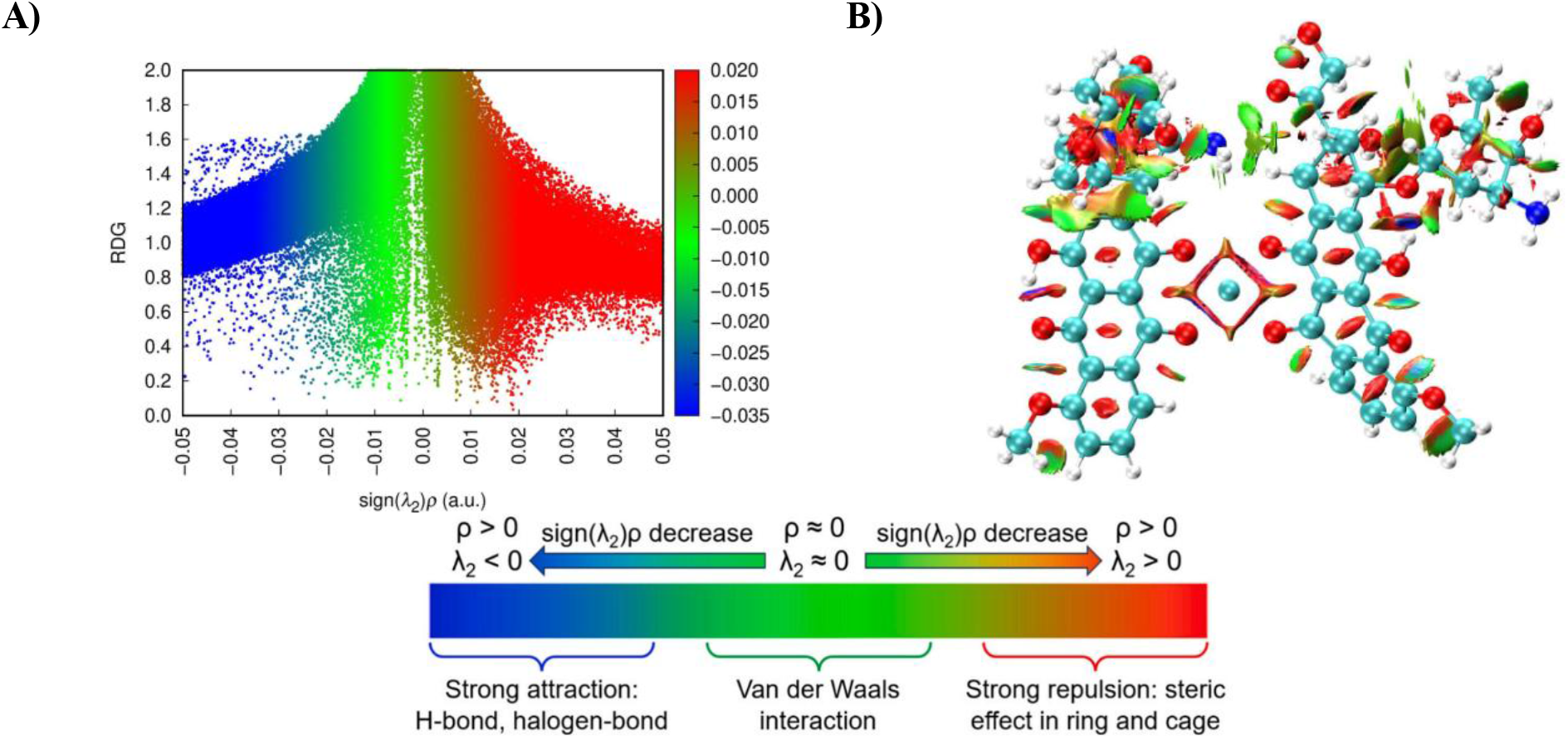
A) RDG scatter plot and B) NCI isosurface of the optimized Cu-DOX complex, illustrating the nature and spatial distribution of its noncovalent interactions. The interaction regions are characterized by sign(λ₂)ρ, where blue, green, and red features denote attractive, weak Van der Waals, and repulsive/steric interactions, respectively.

Thus, compared with free DOX, the most notable change is the appearance of a distinct NCI interaction domain surrounding the Cu²⁺ center, superimposed on the pre-existing dispersive and steric interactions of DOX, supporting the formation of a stable Cu(II)-chelated structure.

#### 3.4.2 Comparative MESP Analysis

Compared with free DOX, the MESP surface of the Cu–DOX complex shows a pronounced redistribution of electrostatic potential around the Cu(II) coordination region and the oxygen-rich anthracycline moiety (Fig. 9). The electron-rich regions, represented by red-to-orange features, remain predominantly localized around the oxygen-containing functionalities, particularly the carbonyl and hydroxyl groups involved in coordination. These regions become spatially reorganized around the Cu(II) center, reflecting the altered electrostatic environment produced by metal chelation. In contrast, the Cu(II) coordination region is surrounded by cyan-to-green potential, indicating a comparatively electron-deficient/intermediate electrostatic environment relative to the oxygen donor sites. The anthracycline framework retains its heterogeneous electrostatic character. At the same time, distinct red/orange regions around the coordinated oxygen functionalities and the corresponding cyan/green region between the donor sites highlight the electrostatic polarization generated upon Cu(II) binding. The amino-sugar moiety also retains localized electrostatic features around its nitrogen- and oxygen-containing groups, although their spatial distribution differs from that of free DOX. Thus, Cu(II) coordination substantially reorganizes the MESP around the oxygen donor sites while retaining the characteristic electrostatic heterogeneity of DOX, providing evidence for a modified electronic environment associated with metal chelation. The observed redistribution of electrostatic potential upon Cu(II) coordination complements the NCI analysis, indicating that metal chelation alters both the noncovalent interaction pattern and the electrostatic environment of DOX.

**Fig. 9.**
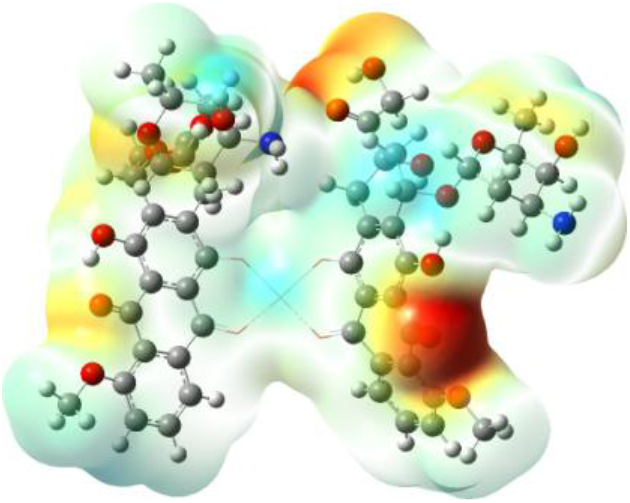
Molecular electrostatic potential (MESP) surface of the DFT-optimized Cu-DOX, illustrating the spatial distribution of electrostatic potential across the anthracycline and amino-sugar moieties. Electron-rich and electron-deficient regions are represented by negative and positive electrostatic potential, respectively, with intermediate colors corresponding to regions of moderate potential.

#### 3.4.3 Comparative IR analysis of free DOX and DOX-Cu complex

The DFT-calculated IR spectra of free DOX and the DOX–Cu complex (Fig. S2) reveal significant changes in vibrational frequencies and intensities upon metal coordination, providing valuable insight into the binding interactions between the ligand and the copper center[17,58,59]. In the spectrum of free DOX, a strong absorption band at 3603 cm⁻¹ is attributed to O–H stretching vibrations arising from the multiple hydroxyl groups present in the anthracycline and daunosamine moieties. Intense bands observed at 1716 and 1668 cm⁻¹ correspond to quinone carbonyl (C=O) stretching vibrations, while the prominent peak at 1589 cm⁻¹ is assigned to aromatic C=C stretching within the conjugated anthraquinone framework. Strong absorptions dominate the fingerprint region at 1299, 1265, 1225, 1125, and 1012 cm⁻¹, originating from C–O, C–O–C, and glycosidic stretching vibrations associated with phenolic, alcoholic, and sugar functionalities. Additionally, the intense band at 940 cm⁻¹ is characteristic of coupled ring-breathing and C–O vibrational modes of the anthracycline skeleton (Table S3) [2,17,58,59].

Upon complexation with Cu, notable spectral changes are observed. The carbonyl stretching band shifts from 1716 cm⁻¹ in free DOX to approximately 1721 cm⁻¹ in the DOX–Cu complex, while additional intense bands appear at 1689, 1639, 1598, and 1574 cm⁻¹. These changes indicate perturbation of the quinone carbonyl groups and enhanced electronic delocalization within the aromatic framework due to metal coordination [17,59]. The oxygen-containing vibrational region also exhibits significant modifications, with strong absorptions at 1336, 1279, 1221, 1189, 1115, 1082, and 1049 cm⁻¹ corresponding to C–O and C–O–C stretching modes. Compared with free DOX, the shifts and increased intensities of these bands suggest the involvement of hydroxyl, carbonyl, and ether oxygen atoms in coordination with the Cu ion. Furthermore, bands at 992 and 975 cm⁻¹ are assigned to ring-breathing and skeletal deformation modes of the anthracycline framework, indicating structural reorganization of the ligand upon complex formation. The low-frequency region below 600 cm⁻¹ becomes particularly important in the metal complex, as it contains skeletal deformation modes and possible Cu–O stretching vibrations that are absent in the spectrum of the free ligand (Table S3) [60].

Overall, the comparison of the two spectra (Table 1) demonstrates that Cu coordination significantly influences the electronic environment of the carbonyl and oxygen-containing functional groups of DOX. The observed shifts in the C=O stretching region, alterations in the C–O/C–O–C vibrational bands, and the emergence of metal–ligand skeletal modes strongly support the formation of a stable DOX–Cu complex. These spectral changes suggest that copper coordination occurs primarily through oxygen donor atoms, particularly the quinone carbonyl and neighboring hydroxyl functionalities, thereby enhancing electron delocalization and stabilizing the complex structure.

**Table 1.** DFT-calculated IR vibrational assignments of free DOX and the DOX–Cu complex, highlighting characteristic vibrational modes and their changes upon Cu(II) coordination.

| Frequency (cm<sup>−1</sup>) | Free DOX | DOX–Cu |
| --- | --- | --- |
| 1716/1721 | Quinone C=O stretch | Coordinated C=O stretch |
| 1668/1689 | Conjugated C=O | Perturbed conjugated C=O |
| 1589/1598 | Aromatic C=C | Metal-influenced aromatic vibration |
| 1299–1225/1336–1221 | C–O, C–O–C | Coordination-affected C–O |
| 940/992 | Ring breathing | Modified skeletal vibration |

#### 3.4.4 Comparative DFT-Raman analysis of free DOX and DOX-Cu complex

The DFT-calculated Raman spectra of free DOX and the DOX–Cu complex reveal substantial changes in vibrational behavior upon metal coordination, providing complementary evidence to the infrared analysis (Fig. S3). Free DOX exhibits Raman activity dominated by the anthraquinone π-conjugated framework (Table S2), with the most intense scattering features located in the 1500–1700 cm⁻¹ region and assigned primarily to conjugated carbonyl and aromatic C=C stretching vibrations. These modes arise from large changes in molecular polarizability associated with vibrations of the extended aromatic system. Following coordination with copper, a pronounced redistribution of Raman intensity is observed across the spectrum, accompanied by significant frequency shifts in several characteristic vibrational modes (Table S3). The most notable change is the dramatic enhancement of Raman scattering in the 1300–1600 cm⁻¹ region, reflecting increased electron delocalization and stronger electronic coupling between the anthraquinone chromophore and the metal center. Vibrational modes associated with oxygen-containing functional groups exhibit substantial shifts and increased intensities, indicating the direct involvement of carbonyl and hydroxyl oxygen atoms in coordination.

Furthermore, the appearance of low-frequency Raman-active modes assigned to Cu–O stretching vibrations provides direct spectroscopic evidence for metal–ligand bond formation. The observed spectral modifications demonstrate that copper coordination affects not only the local donor atom environment but also the overall electronic structure of the anthracycline framework. The Raman response undergoes substantial redistribution upon Cu(II) coordination, with several modes exhibiting pronounced frequency shifts and changes in relative intensities, reflecting the mixing of carbonyl, aromatic, and C–O vibrational modes (Table 2) [15,58,61,62]. The combined Raman and IR results therefore support a coordination model in which copper interacts primarily with the quinone carbonyl, with localization, altered molecular polarizability, and greater structural stabilization of the resulting DOX–Cu complex.

**Table 2.**
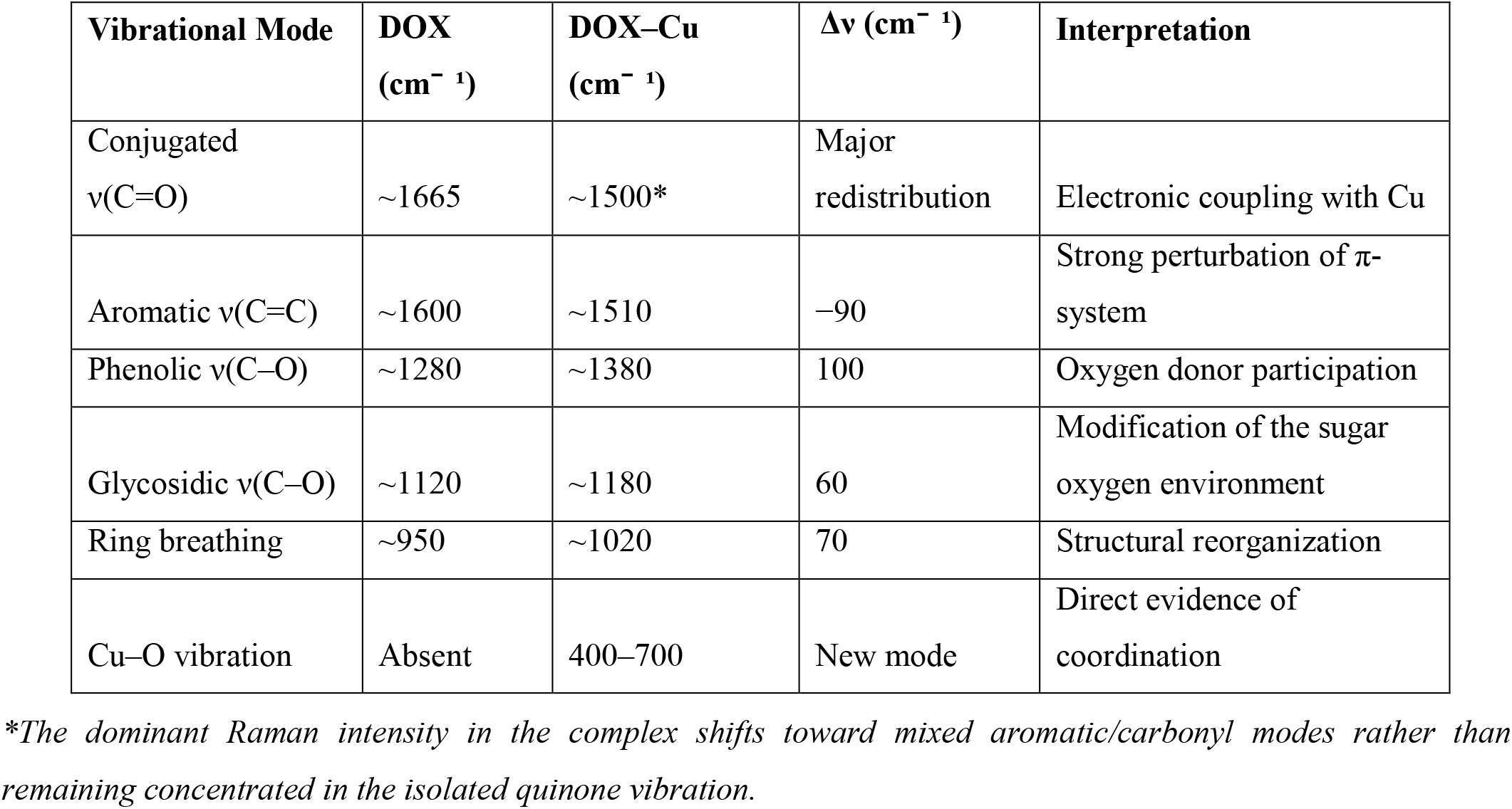
Comparative Raman frequency shifts between free DOX and the DOX–Cu complex, highlighting the vibrational changes associated with Cu(II) coordination and the resulting structural and electronic perturbations.

| Vibrational Mode | DOX<br>(cm<sup>−1</sup>) | DOX–Cu<br>(cm<sup>−1</sup>) | $\Delta\nu$ (cm<sup>−1</sup>) | Interpretation |
| --- | --- | --- | --- | --- |
| Conjugated<br>$\nu(\text{C}=\text{O})$ | ~1665 | ~1500* | Major<br>redistribution | Electronic coupling with Cu |
| Aromatic $\nu(\text{C}=\text{C})$ | ~1600 | ~1510 | −90 | Strong perturbation of π-<br>system |
| Phenolic $\nu(\text{C}-\text{O})$ | ~1280 | ~1380 | 100 | Oxygen donor participation |
| Glycosidic $\nu(\text{C}-\text{O})$ | ~1120 | ~1180 | 60 | Modification of the sugar<br>oxygen environment |
| Ring breathing | ~950 | ~1020 | 70 | Structural reorganization |
| Cu–O vibration | Absent | 400–700 | New mode | Direct evidence of<br>coordination |

*\*The dominant Raman intensity in the complex shifts toward mixed aromatic/carbonyl modes rather than remaining concentrated in the isolated quinone vibration.*

**Table 3.** Binding constants (*K*) determined from UV–Vis spectroscopic studies.

| System | Binding constant (K) $\text{M}^{-1}$ | Gibbs free energy change ($\Delta G$ at 298.15 K) $\text{kJ mol}^{-1}$ |
| --- | --- | --- |
| DOX–Cu(II) | $1.626 \times 10^4$ | –24.04 |
| EB - DNA | $2.2468 \times 10^4$ | –24.84 |
| DOX - DNA | $2.059 \times 10^4$ | –24.62 |
| DOX - Cu(II) - DNA | $20.561 \times 10^4$ | –30.33 |

**Table 4.** Fluorescence-derived DNA-binding parameters of EB, DOX, and Cu(II)–DOX.

| System | Stern-Volmer (SV) analysis |  | Modified SV analysis |  | Double log plot constants |  | Hill binding analysis |  |  |
| --- | --- | --- | --- | --- | --- | --- | --- | --- | --- |
| | $K_{SV} (M^{-1})$ | $K_q (M^{-1} s^{-1})$ | $f_a$ | $K_a (M^{-1})$ | $K (M^{-1})$ | $n$ | $n_H$ | $K_H (M^{-1})$ | $EC_{50}/IC_{50} (\mu M)$ |
| EB-DNA | - | - | - | - | $2.13 \times 10^4$ | 1.07 | 1.67 | $2.88 \times 10^4$ | $EC_{50} = 34.77$ |
| EB- DNA-<br>DOX | $1.166 \times 10^5$ | $5.83 \times 10^{12}$ | 0.686 | $2.41 \times 10^5$ | $1.15 \times 10^4$ | 0.81 | 1.48 | $3.87 \times 10^5$ | $IC_{50} = 2.59$ |
| EB -DNA -<br>Cu(II)- DOX | $1.10 \times 10^5$ | $5.58 \times 10^{12}$ | 0.535 | $1.26 \times 10^6$ | $4.46 \times 10^2$ | 0.51 | 0.55 | $2.39 \times 10^5$ | $IC_{50} = 4.18$ |

Thus far, the computational results establish two key features: (i) stable intercalative binding of DOX with DNA and (ii) Cu(II) chelation that alters the structural and electronic characteristics of DOX. Complexation with Cu(II) reduces the accessible degrees of conformational freedom of DOX. It introduces additional structural rigidity, which may favor a more defined and intercalation-compatible orientation of the anthracycline moiety within the DNA base-pair stack. The resulting structural restriction may reduce the conformational penalty associated with DNA binding, while the coordinated Cu(II) center could provide additional stabilization through interactions with DNA donor sites. Additionally, the Cu(II) center may facilitate secondary stabilization through metal-mediated interactions with nucleobase donor atoms or phosphate oxygen atoms within the DNA framework [10]. However, conventional molecular docking methods are generally limited in their ability to accurately describe transition-metal complexes because empirical force fields may not adequately represent metal coordination environments [9]. Therefore, experimental studies were undertaken to evaluate the influence of Cu(II) on DOX–DNA binding. UV– Vis absorption and fluorescence spectroscopy were employed to quantitatively characterize the binding behavior and assess the role of Cu(II) in the DOX–DNA interaction.

### 3.5 UV–Vis Spectroscopic Studies

Electronic absorption spectroscopy is a widely employed technique for investigating drug–DNA interactions [2]. Association of small molecules with DNA can produce hypochromism, broadening of the absorption envelope, and shifts in the absorption maxima. Such spectral changes are particularly characteristic of intercalative interactions, arising from the close association of the ligand chromophore with DNA base pairs [2,63]. EB, a classical planar DNA intercalator, was therefore investigated as a reference system to establish the characteristic spectroscopic response associated with DNA binding (Fig. S4) [45,63]. The UV–Vis studies of DOX and the DOX–Cu(II) system were subsequently conducted to assess their DNA-binding behavior experimentally and corroborate the computational results.

The interaction of DOX with herring sperm DNA was investigated by maintaining the DOX concentration at 80 μM while varying the DNA concentration from 0 to 136.4 μM. Free DOX exhibited a characteristic absorption band in the 450–550 nm region, with an absorption maximum at 483 nm. The absorption of DOX in this region has previously been reported near 488 nm and assigned to a π→π* transition involving the carbonyl and C–OH groups [21,64]. Upon addition of DNA, the absorption maximum of DOX at 483 nm exhibited approximately 50% hypochromism accompanied by a bathochromic shift of approximately 17 nm at a DOX molar ratio of 1:1.1 (Fig. 10A). The pronounced hypochromism together with the appreciable bathochromic shift is characteristic of close interaction between the aromatic chromophore and DNA base pairs and has been associated with intercalative binding [2,63,64]. The substantial decrease in DOX absorptivity, accompanied by the red shift, therefore indicates a strong association of the DOX chromophore with DNA, consistent with an intercalative mode of binding. The binding constant obtained from the double-reciprocal analysis was 2.059 x 10^4^ M^-1^ (Equ 2.1, Fig. S5), which is comparable to that of EB (K = 2.247 x 10^4^ M^-1^). The spectroscopic observations shown in Fig. 10A agree with the molecular docking and molecular dynamics simulation results presented earlier, which demonstrated a stable association between DOX and the DNA helix. Thus, the computational and spectroscopic results provide mutually supportive evidence for strong intercalative binding of DOX to DNA.

**Fig. 10.**
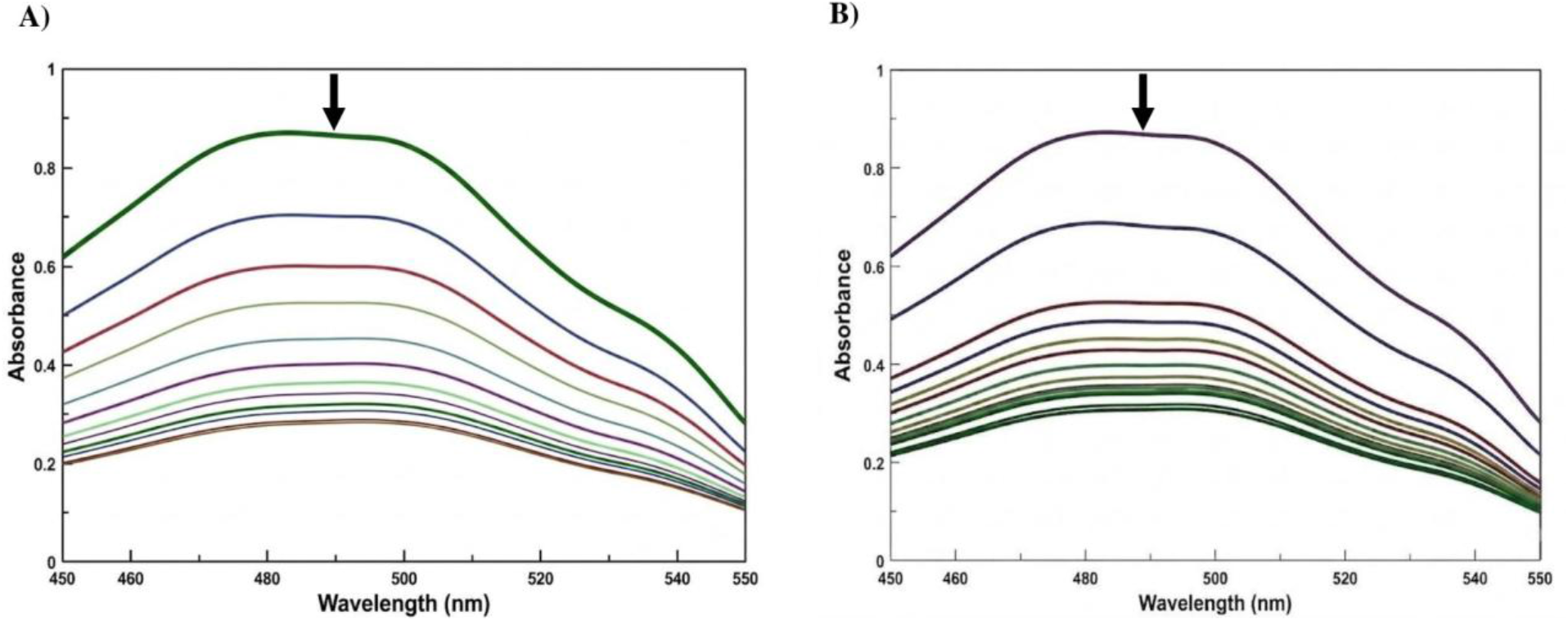
Spectrophotometric titration of DOX and DOX-Cu(II) complex in the presence of DNA in a 10 mM phosphate buffer of pH 7.0 at room temperature. A) Curves (from top to bottom) denote the titration of DOX (80 μM) treated with 0, 12.4,24.8, 37.2, 49.6, 62, 74.4, 86.8, 99.2, 111.6, 124, and 136.4 μM of DNA. B) Curves (from top to bottom) denote the titration of DOX-Cu(II) complex (1:0.5) treated with 0, 6.2, 12.4, 18.6, 24.8, 31, 37.2, 43.4, 49.6, 55.8, 62, 68.2, and 74.4 μM of DNA.

The interaction of DOX with Cu(II) was investigated by titrating a fixed concentration of DOX (80 μM) with increasing concentrations of CuCl_2_ over the range 0 –72 μM. With increasing Cu(II) concentration, the absorption maximum of DOX at 484 nm decreased by more than 50%. It underwent a bathochromic shift to 497 nm, indicating a pronounced interaction between DOX and Cu(II) (Fig. S6A). Similar spectral changes have been reported for the complexation of other bioactive ligands with Cu(II) ions[17,19,59]. The binding constant obtained from the double-reciprocal analysis was 1.626 x 10^4^ M^-1^, with the corresponding plot showing excellent linearity (r^2^ > 0.98), indicating an appreciable interaction between DOX and Cu(II) (Fig. S6 B). The spectral response approached saturation around a DOX–Cu(II) molar ratio of 1:0.5, indicating progressively smaller spectral changes upon further addition of Cu(II). Therefore, a DOX–Cu(II) molar ratio of 1:0.5 was selected for subsequent DNA-binding studies, as it represents a Cu(II) concentration below the near-saturation region while retaining the DOX–Cu(II) system’s pronounced spectroscopic response. These experimental findings complement the computational results presented earlier, which collectively support the formation and structural characteristics of the DOX–Cu(II) complex.

The DNA-binding behavior of the DOX–Cu(II) system was subsequently investigated at a fixed DOX–Cu(II) molar ratio of 1:0.5 while the DNA concentration was progressively increased. At a DOX–Cu(II) molar ratio of 1:0.5:0.6, the absorption maximum of DOX at 483 nm exhibited approximately 60% hypochromism accompanied by a bathochromic shift of approximately 12 nm (Fig. 10B). The substantial decrease in absorptivity, together with the pronounced red shift, indicates a strong interaction between the DOX–Cu(II) system and DNA. Such spectral changes are characteristic of the close association of an aromatic chromophore with DNA base pairs and are consistent with an intercalative mode of interaction [2,63,64]. The binding constant obtained from the double-reciprocal analysis was 20.561 x 10^4^ M^-1^ (Fig. S7). Thus, relative to the DOX–DNA system, incorporation of Cu(II) produces an approximately 10-fold increase in the apparent DNA association constant.

The standard free-energy change at room temperature T = 298.15 K for the EB–DNA system was approximately −24.8 kJ mol⁻¹, while that for the DOX–DNA system was approximately −24.6 kJ mol^-1^ (Equ. 2.2). The comparable negative ΔG values indicate that the interaction of DOX with DNA is thermodynamically favourable and of comparable stability to the well-established EB–DNA interaction. The corresponding standard free-energy change becomes substantially more favourable, from −24.6 kJ mol^-1^ for DOX–DNA to −30.3 kJ mol^-1^ for the DOX–Cu(II)–DNA system. Thus, incorporation of Cu(II) results in an additional free-energy stabilization of approximately 5.7 kJ mol^-1^ in the ternary DOX–Cu(II)–DNA system relative to the DOX–DNA system. This enhanced thermodynamic stabilization is consistent with the approximately tenfold increase in the association constant for DOX–DNA in the presence of Cu(II) ions. These results demonstrate that Cu(II) coordination substantially enhances the thermodynamic stabilization of the DNA-associated DOX system. Taken together with the observed hypochromism and bathochromic shift, the spectroscopic data support intercalative association of the DOX chromophore with DNA and further indicate that Cu(II) coordination strengthens this DNA-associated interaction. Overall, the UV–Vis results demonstrate that DOX binds favourably to DNA, with Cu(II) coordination producing a marked enhancement in both the association constant and thermodynamic stability of the resulting ternary complex.

### 3.6 Fluorescence Spectroscopic Studies

Fluorescence spectroscopy provides a sensitive approach for probing ligand–DNA interactions, particularly when a fluorescent DNA-binding probe is employed [20,46,63]. In the present study, EB was employed as a fluorescent reporter to monitor DOX interaction with DNA via competitive displacement of DNA-bound EB. The EB–DNA system was therefore first established under the experimental conditions used for the competitive studies and was subsequently considered as the reference system for evaluating the effects of DOX and Cu(II)-coordinated DOX.

EB was maintained at a fixed concentration of 15 μM throughout the fluorescence experiments. Free EB exhibited a weak emission band between 550 and 650 nm, with a maximum at approximately 596 nm upon excitation at 480 nm. Progressive addition of DNA resulted in a pronounced enhancement of the EB emission without a substantial displacement of the emission maximum (Fig. S8A) [41,46,63].

The fluorescence intensity approached a plateau at an EB-DNA molar ratio of approximately 1:6.6, indicating that the majority of the EB molecules had become associated with DNA under the experimental conditions. The EB-DNA ratio of 1:4 was selected for the competitive experiments because the fluorescence intensity was approximately twice that of free EB. The competitive binding experiments were therefore performed using the EB– DNA system maintained at this fixed molar ratio. DOX itself exhibits comparatively weak fluorescence under the experimental aqueous conditions; therefore, changes in the emission of DNA-bound EB provide an indirect means of monitoring the interaction of DOX with DNA. Upon progressive addition of DOX, the emission intensity of EB–DNA at approximately 593 nm decreased systematically, with only a minor change in the emission maximum (Fig. 11A). The decrease in EB fluorescence reached an approximately saturated region at higher DOX concentrations, demonstrating concentration-dependent perturbation and displacement of DNA-bound EB. This behaviour is consistent with t h e competitive occupation of DNA-associated sites by DOX. Similar EB-based competitive fluorescence approaches are widely used as indirect probes of DNA binding by small molecules and metal complexes [65–68].

**Fig. 11.**
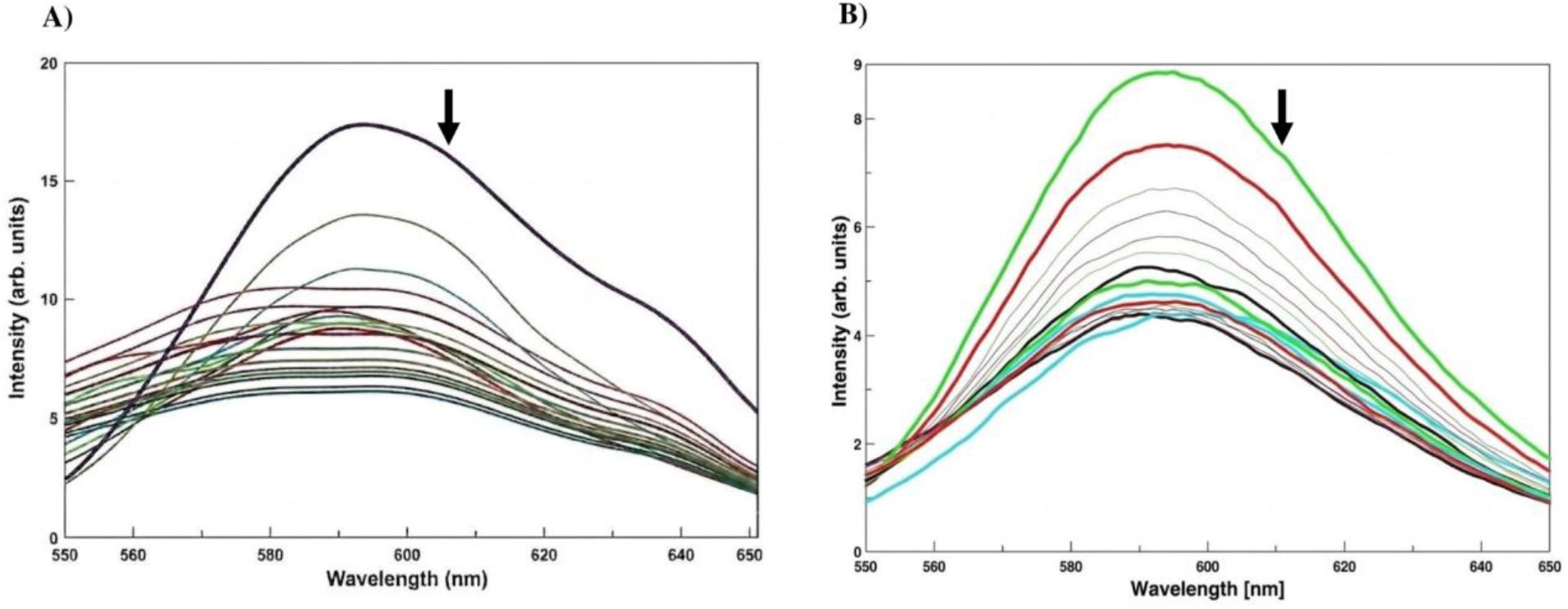
Spectrofluorimetric titration of EB-DNA and EB-DNA-Cu(II) complex in the presence of DOX in a 10 mM CP buffer of pH 7.0 at room temperature. A) Curves (from top to bottom) denote the titration of EB-DNA (1:0.4) treated with DOX 0, 2, 4, 6, 8, 10, and 12 μM of DOX. B) Curves (from top to bottom) denote the titration of EB-DNA-Cu(II) complex (1:0.4:1.2) treated with 0, 9.12, 6.49, 5.85, 5.79, 5.59, 5.39, 5.11, 5.06, 4.96, 4.72, 4.63, 4.58, and 4.48 μM of DOX.

As the reference system, the EB–DNA interaction was quantitatively evaluated using the same fluorescence models subsequently applied to the competitive systems. The conventional double-log treatment yielded a binding parameter of K = 2.13 x 10^4^ M^-1^ with n = 1.07 (Fig. S8B). The corresponding nonlinear Hill analysis gave K_H_ = 2.88 x 10^4^ M^-1^, n_H_ = 1.67, (r^2^>0.99), and an EC_50_ value (half-maximal effective concentration: the concentration of the DNA producing 50% enhancement of the maximum fluorescence intensity) of approximately 34.77 μM, (Fig. S8C). The value of n_H_ > 1 indicates positive apparent cooperativity in the EB fluorescence response to DNA. However, the Hill coefficient should be regarded as an empirical descriptor of the overall binding response rather than as direct evidence of a specific molecular stoichiometry. These parameters establish the characteristic fluorescence response of the EB–DNA reference system and provide the basis for assessing the changes produced by DOX and Cu(II)-coordinated DOX. Since DNA addition enhances rather than quenches EB fluorescence in the reference EB–DNA system, conventional SV and modified SV parameters (K_SV_, K_q,_ K_a_, and f_a_), which describe fluorescence quenching, are not applicable and are therefore not included for this system.

The fluorescence quenching data were initially evaluated using the SV relationship (Eq. 2.3). For the EB–DNA– DOX system, the linear Stern–Volmer analysis gave K_SV_ = 1.166 x 10^5^ M^-1^, corresponding to an apparent bimolecular quenching constant, K_q_ of 5.83 x 10^12^ M^-1^s^-1^ when τ_o_ = 2 x 10^-8^ s is used (Fig. S9 A). The magnitude of K_q_, which is considerably higher than the diffusion-controlled range expected for purely collisional quenching, indicates that the observed fluorescence attenuation is consistent with a substantial contribution from static association and displacement of EB from DNA.

The modified Stern–Volmer treatment provided additional information on the accessibility of the DNA-associated EB population (Eq. 2.5, Fig. S9B). For EB–DNA–DOX, the accessible fraction was f_a_ = 0.686, indicating that approximately 68.6% of the initial EB fluorescence was accessible to DOX-induced quenching. At the same time, the corresponding apparent association parameter was K_a_ = 2.41 × 10^4^ M^-1^. The f_a_ value indicates that the fluorescence response does not represent an entirely homogeneous EB population with identical accessibility to DOX. Importantly, K_a_ obtained from the modified SV treatment is an association parameter for the accessible population and should not be regarded as the overall thermodynamic DOX–DNA binding constant. The nonlinear Hill treatment further characterized the concentration dependence of the competitive displacement process (Equ 2.6, Fig. S9C). For EB–DNA–DOX, the complete quenching profile yielded KH = 3.87 × 10^5^ M^-1^, IC_50_ = 2.59 μM (IC₅₀ is the half-maximal inhibitory concentration: the DOX concentration producing 50% inhibition of the maximum fluorescence intensity), and n_H_ = 1.48, with an excellent fit (r²> 0.99). The IC_50_ value is markedly lower than the approximately 34.77 μM obtained for the EB–DNA reference response, demonstrating the strong concentration-dependent effect of DOX on the DNA-associated EB system. The decrease in n_H_ from 1.67 for EB– DNA to 1.48 for EB–DNA–DOX indicates a change in the apparent cooperative character of the fluorescence response. In contrast, the value remaining above unity indicates positive apparent cooperativity in the displacement process. The Hill parameters, therefore, provide an independent description of the DOX-induced change in the EB–DNA competitive system.

The conventional double-log analysis provides particularly useful quantitative evidence for the competitive displacement interpretation (Fig. S9D). For EB–DNA–DOX, the apparent binding parameter was K = 1.15 x 10^4^ M^-1^, with n = 0.807, compared with K = 2.13 x 10^4^ M^-1^ and n = 1.07 for the EB–DNA reference system (Fig. S9D). The reduction in K should not be interpreted as a lower intrinsic binding constant of DOX for DNA. In the present fluorescence experiment, the double-log parameter describes the response of the pre-existing EB–DNA system to the addition of the competing DOX molecule. Therefore, the decrease in K, together with the decrease in n, represents a quantitative change in the competitive EB–DNA system produced by DOX and is consistent with effective competition between DOX and EB for DNA-associated binding sites. The fluorescence-derived K is consequently a model-dependent competitive parameter and is distinct from the thermodynamic association constant obtained from the independent UV–Vis absorption measurements.

The effect of Cu(II) coordination on the DOX-induced displacement of DNA-bound EB was subsequently examined using a fixed EB-DNA-Cu(II) molar ratio of 1:4:0.4, followed by progressive addition of DOX. Addition of DOX produced a pronounced decrease in EB fluorescence with only a minor change in the emission maximum, demonstrating substantial displacement of the DNA-associated EB population (Fig. 11 B). Approximately 50% attenuation of the initial EB fluorescence occurred at a DOX concentration of approximately 4–5 μM, after which the fluorescence response approached a plateau. Notably, the initial EB fluorescence in the Cu(II)-containing system was approximately 8.87 a.u., compared with 16.15 a.u. for the corresponding EB– DNA system in the absence of Cu(II), representing an approximately 45% reduction before DOX addition. Thus, Cu(II) itself produces a substantial change in the EB–DNA reference response before DOX is introduced. The subsequent fluorescence titration, therefore, represents DOX-induced competition within a Cu(II)-containing DNA system rather than a simple repetition of the EB–DNA–DOX experiment.

The SV analysis of the EB–DNA–Cu(II)–DOX system gave K_SV_ = 1.10 x 10^5^ M^-1^, with a corresponding K_q_ =5.58 x 10^12^ M^-1^s^-1^ (Fig. S10 A). The K_q_ value is comparable to that obtained for the EB–DNA–DOX system, indicating static ground state quenching in the presence of Cu(II), which could be attributed to the competitive interaction of DOX with the DNA-associated EB system. The modified SV analysis showed a further pronounced change in the accessible fluorescence population. The value of f_a_ decreased from 0.686 for EB–DNA–DOX to 0.535 for EB– DNA–Cu(II)–DOX, indicating that the fraction of the EB population accessible to DOX-induced quenching was substantially reduced in the presence of Cu(II) (Fig. S10 B). At the same time, the apparent K_a_ increased from 2.41 x 10^5^ M^-1^ to 1.26 x 10^6^ M^-1^, corresponding to an approximately 5.2-fold increase. This is consistent with enhanced occupation or perturbation of DNA binding sites by the Cu–DOX system. Importantly, the reduced f_a_ reflects the accessibility of the EB reporter population rather than a decrease in the intrinsic DOX–DNA affinity.

The Hill analysis likewise demonstrated a substantial change in the fluorescence displacement behaviour. For EB– DNA–Cu(II)–DOX, the fitted values were K_H_ = 2.39 x 10^5^ M^-1^, IC_50_ = 4.18 μM, and n_H_ = 0.55, with r^2^ > 0.99 (Fig. S10C). Relative to the EB–DNA reference, the Hill coefficient decreases from 1.67 to 0.55 through the DOX-containing system (n_H_=1.48), indicating a pronounced change in the cooperative character of the fluorescence response. The sub-unity value in the Cu-containing system is consistent with increased heterogeneity and/or non-equivalent accessibility of DNA-associated sites in the competitive displacement process. The increase in IC_50_ from 2.59 μM for EB–DNA–DOX to 4.18 μM for EB–DNA–Cu(II)–DOX indicates that the concentration dependence of EB displacement is substantially different after Cu(II) coordination. These Hill parameters therefore demonstrate that Cu(II) reorganizes the competitive displacement behaviour rather than simply producing a uniform increase in fluorescence quenching. The conventional double-log treatment provides a clear quantitative comparison of the competitive fluorescence response across the three systems. The K decreases progressively from 2.13 x 10^4^ M^-1^ for the EB–DNA reference to 1.15 x 10^4^ M^-1^ for EB–DNA–DOX and further to 4.46 x 10^2^ M^-1^ for EB–DNA–Cu(II)–DOX (Fig. S10D).

Thus, compared with the EB–DNA reference, the K value decreases by approximately 46% in the presence of DOX and by approximately 98% in the presence of Cu(II)-coordinated DOX. The corresponding n values decrease from 1.07 to 0.81, then to approximately 0.51. Because these parameters are obtained from the competitive fluorescence response of DNA-bound EB, the progressive decrease in K should be interpreted as a quantitative indication of increasingly pronounced competition and effective displacement of EB. The particularly large reduction in K in the Cu(II)-containing system therefore provides strong fluorescence evidence that the interaction of DOX with DNA is substantially enhanced upon Cu(II) coordination, resulting in a much stronger perturbation of the pre-existing EB–DNA binding equilibrium.

The agreement between the fluorescence and UV–Vis results is particularly significant because the two techniques probe the systems from different perspectives. The fluorescence experiment, using DNA-bound EB as the reference, independently shows that the apparent competitive K of the EB system decreases dramatically when DOX is introduced and decreases even more strongly when DOX is coordinated with Cu(II). This progressive reduction in the EB competitive parameter is therefore consistent with increasingly effective competition by DOX for DNA-associated binding sites, with the strongest competition occurring for the Cu(II)-coordinated DOX system.

### 3.7 Future Scope

The present study offers a molecular and spectroscopic foundation to better understand the role of Cu(II) coordination in modifying the structure and DNA-binding behavior of DOX. However, further computational studies are required to extend these observations to the complete DOX–Cu(II)–DNA system. Molecular dynamics simulations of the ternary complex, in particular, would give valuable atomistic insight into the stability of the metal-chelated drug in the DNA intercalation site, the persistence of Cu–O coordination and possible secondary interactions of the metal center with DNA bases or phosphate groups. This would complement the existing 100 ns MD study of DOX-DNA and help establish if the experimentally reported improvement in DNA binding is maintained dynamically in the presence of Cu(II).

Apart from the study at the molecular level, the biological significance of the Cu-mediated modulation of DOX has to be empirically demonstrated. In metal-loading investigations, the stoichiometry, loading efficiency and stability of the DOX–Cu complex should be determined first in physiologically relevant conditions. This would give quantitative confirmation of how much Cu(II) is linked with DOX and how persistent the complex is in biological media. Cellular studies comparing free DOX and DOX–Cu would be necessary to see if the changed DNA binding translates into differences in cellular uptake, cytotoxicity and anticancer activity. Finally, bioavailability and pharmacokinetic studies of the metal-chelated medication will be necessary in an effort to determine if Cu(II) coordination affects the absorption, stability, distribution and persistence of DOX in biological systems. In summary, these studies would link the existing spectroscopic and computational data to their potential biological and therapeutic relevance, leading to a more robust assessment of whether Cu(II) coordination can be used as a viable strategy to modulate the activity of DOX

## 4. Conclusion

In conclusion, this study demonstrates that Cu(II) coordination substantially modulates the structural, electronic, and DNA-binding properties of doxorubicin (DOX), as established through complementary computational and spectroscopic approaches. Molecular docking identified a favorable intercalative binding mode of DOX within DNA, with a binding affinity of −7.84 kcal mol⁻ ¹. Molecular dynamics simulations of 100 ns duration confirmed the stability of the resulting complex, with no significant dissociation. DFT calculations indicated coordination of Cu(II) through the oxygen-rich region of the anthracycline framework, accompanied by distinct changes in the local structural and electronic environment of DOX. These findings were further supported by NCI, MESP, IR, and Raman analyses, which collectively indicated increased structural rigidity and reduced conformational freedom following Cu(II) coordination.

Experimental investigations demonstrated that Cu(II) coordination resulted in a pronounced enhancement of DOX–DNA association. The experimentally determined binding constant increased from approximately 2.059 × 10⁴ M⁻ ¹ for DOX–DNA to 20.561 × 10⁴ M⁻ ¹ for DOX–Cu(II)–DNA, corresponding to an approximately ten fold increase in the apparent DNA-binding affinity. The more favorable Gibbs free energy further supported the enhanced thermodynamic stability of the Cu(II)-coordinated complex. Fluorescence displacement studies further demonstrated that Cu(II)–DOX more effectively perturbed the pre-existing EB–DNA equilibrium than DOX alone, consistent with a static quenching mechanism. This enhanced competitive behavior was accompanied by an approximately 5.2-fold increase in the modified Stern–Volmer association constant (Kₐ), from 2.41 × 10⁵ M⁻ ¹ for DOX to 1.26 × 10⁶ M⁻ ¹ for Cu(II)–DOX. The associated changes in the competitive binding parameters further suggested that Cu(II) coordination alters the apparent mode and heterogeneity of DNA-site occupation.

These findings indicate that Cu(II) coordination induces structural and electronic modifications in DOX that substantially alter its interaction with DNA and enhance its DNA-binding affinity. The integration of computational and experimental evidence provides a mechanistic basis for understanding how metal coordination can modulate the biomolecular recognition properties of anthracycline drugs. Further metal-aware computational studies and biological and pharmacological investigations are required to determine whether the enhanced DNA association of Cu–DOX translates into altered anticancer activity and therapeutic potential.

## Acknowledgement

The authors gratefully acknowledge Bhagawan Sri Sathya Sai Baba, the Founder Chancellor of SSSIHL, for His constant guidance. The authors also thank the administration of SSSIHL for providing SAI-HPC, the Central Research Instruments Facility (CRIF), and research support. Dr. Aditya acknowledges the support from DST through DST INSPIRE Faculty Fellowship.

## Declaration of competing interest

There are no relevant financial or non-financial interests to disclose.

## Declaration of generative AI use

During the preparation of this work, the authors used OpenAI’s ChatGPT (GPT-5.6 Luna) to assist i n refining the draft into an academic style. Following its use, the authors reviewed and edited the content as necessary and took full responsibility for the published article.

**Supplementary materials** are available.

## Credit authorship contribution statement

**Dipanshu Ranjan Pattanayak:** Validation, Formal Analysis, Investigation, Data Curation, Writing – Original Draft, Visualization, **R Dharmaraj:** Conceptualization, Formal Analysis, Investigation, Data Curation, Writing - Original Draft, **R Sarojini:** Conceptualization, Formal Analysis, **Aditya Dileep Kurdekar:** Investigation, Writing-Review & Editing, **Chelli Sai Manohar:** Data Curation, Writing-Review & Editing.

## Notes

### Competing Interest Statement

The authors have declared no competing interest.

## References

[1] A.A. Almaqwashi, T. Paramanathan, I. Rouzina, M.C. Williams, Mechanisms of small molecule– DNA interactions probed by single-molecule force spectroscopy, Nucleic Acids Res. 44 (2016) 3971–3988. 10.1093/nar/gkw237.

[2] D. Agudelo, P. Bourassa, G. Bérubé, H.A. Tajmir-Riahi, Review on the binding of anticancer drug doxorubicin with DNA and tRNA: Structural models and antitumor activity, J. Photochem. Photobiol. B 158 (2016) 274–279. 10.1016/j.jphotobiol.2016.02.032.

[3] G.L. Beretta, F. Zunino, Molecular Mechanisms of Anthracycline Activity, in: K. Krohn (Ed.), Anthracycline Chem. Biol. II, Springer Berlin Heidelberg, Berlin, Heidelberg, 2007: pp. 1–19. 10.1007/128_2007_3.

[4] H. Zhang, Y.G. Gao, G.A. Van Der Marel, J.H. Van Boom, A.H. Wang, Simultaneous incorporations of two anticancer drugs into DNA. The structures of formaldehyde-cross-linked adducts of daunorubicin-d(CG(araC)GCG) and doxorubicin-d(CA(araC)GTG) complexes at high resolution, J. Biol. Chem. 268 (1993) 10095–10101. 10.1016/S0021-9258(18)82176-7.

[5] J.L. Nitiss, Targeting DNA topoisomerase II in cancer chemotherapy, Nat. Rev. Cancer 9 (2009) 338–350. 10.1038/nrc2607.

[6] S.Y. Van Der Zanden, X. Qiao, J. Neefjes, New insights into the activities and toxicities of the old anticancer drug doxorubicin, FEBS J. 288 (2021) 6095–6111. 10.1111/febs.15583.

[7] F.S. Carvalho, A. Burgeiro, R. Garcia, A.J. Moreno, R.A. Carvalho, P.J. Oliveira, Doxorubicin-Induced Cardiotoxicity: From Bioenergetic Failure and Cell Death to Cardiomyopathy, Med. Res. Rev. 34 (2014) 106–135. 10.1002/med.21280.

[8] U. Ndagi, N. Mhlongo, M. Soliman, Metal complexes in cancer therapy – an update from drug design perspective, Drug Des. Devel. Ther. Volume11 (2017) 599–616. 10.2147/DDDT.S119488.

[9] M.J. Hannon, Metal-based anticancer drugs: From a past anchored in platinum chemistry to a post-genomic future of diverse chemistry and biology, Pure Appl. Chem. 79 (2007) 2243–2261. 10.1351/pac200779122243.

[10] P. Liu, Z.C. Han, Treatment of acute promyelocytic Leukemia and other hematologic malignancies with arsenic trioxide: Review of clinical and basic studies, Int. J. Hematol. 78 (2003) 32–39. 10.1007/BF02983237.

[11] C. Marzano, M. Pellei, F. Tisato, C. Santini, Copper Complexes as Anticancer Agents, Anticancer Agents Med. Chem. 9 (2009) 185–211. 10.2174/187152009787313837.

[12] D. Denoyer, S. Masaldan, S. La Fontaine, M.A. Cater, Targeting copper in cancer therapy: ‘Copper That Cancer,’ Metallomics 7 (2015) 1459–1476. 10.1039/C5MT00149H.

[13] S. Bhuiya, S. Chowdhury, L. Haque, S. Das, Spectroscopic, photophysical and theoretical insight into the chelation properties of fisetin with copper (II) in aqueous buffered solutions for calf thymus DNA binding, Int. J. Biol. Macromol. 120 (2018) 1156–1169. 10.1016/j.ijbiomac.2018.08.162.

[14] S. Goswami, S. Ray, M. Sarkar, Spectroscopic studies on the interaction of DNA with the copper complexes of NSAIDs lornoxicam and isoxicam, Int. J. Biol. Macromol. 93 (2016) 47–56. 10.1016/j.ijbiomac.2016.08.025.

[15] P.K. Dutta, J.A. Hutt, Resonance Raman spectroscopic studies of adriamycin and copper(II)-adriamycin and copper(II)-adriamycin-DNA complexes, Biochemistry 25 (1986) 691–695. 10.1021/bi00351a028.

[16] V. Malatesta, A. Gervasini, F. Morazzoni, Chelation of copper(II) ions by doxorubicin and 4′-epidoxorubicin: ESR evidence for a new complex at high anthracycline/copper molar ratios, Inorganica Chim. Acta 136 (1987) 81–85. 10.1016/S0020-1693(00)87099-1.

[17] M. Feng, Y. Yang, P. He, Y. Fang, Spectroscopic studies of copper(II) and iron(II) complexes of adriamycin, Spectrochim. Acta. A. Mol. Biomol. Spectrosc. 56 (2000) 581–587. 10.1016/S1386-1425(99)00157-2.

[18] X. Yuan, D. Guo, M. Zhang, The influence of Cu(II), Mg(II) on the binding of adriamycin with DNA and the study on their interaction mechanism, Spectrochim. Acta. A. Mol. Biomol. Spectrosc. 63 (2006) 444–448. 10.1016/j.saa.2005.05.029.

[19] W. Zhang, P. Zhang, X. Xu, M. Li, S. Wang, H. Mu, K. Sun, Synergy effects of copper ion in doxorubicin-based chelate prodrug for cancer chemo-chemodynamic combination therapy, Drug Deliv. 30 (2023) 2219426. 10.1080/10717544.2023.2219426.

[20] Y. Sun, S. Bi, D. Song, C. Qiao, D. Mu, H. Zhang, Study on the interaction mechanism between DNA and the main active components in Scutellaria baicalensis Georgi, Sens. Actuators B Chem. 129 (2008) 799–810. 10.1016/j.snb.2007.09.082.

[21] S. Gómez, P. Lafiosca, F. Egidi, T. Giovannini, C. Cappelli, UV-Resonance Raman Spectra of Systems in Complex Environments: A Multiscale Modeling Applied to Doxorubicin Intercalated into DNA, J. Chem. Inf. Model. 63 (2023) 1208–1217. 10.1021/acs.jcim.2c01495.

[22] A. Nedělníková, P. Stadlbauer, M. Otyepka, P. Kührová, M. Paloncýová, Atomistic Insights Into Interaction of Doxorubicin With DNA : From Duplex to Nucleosome, J. Comput. Chem. 46 (2025) e70035. 10.1002/jcc.70035.

23. M.J. Frisch, G.W. Trucks, H.B. Schlegel, G.E. Scuseria, M.A. Robb, J.R. Cheeseman, G. Scalmani, Gaussian 16, (2016).

[24] A.D. Becke, Density-functional thermochemistry. III. The role of exact exchange, J. Chem. Phys. 98 (1993) 5648–5652. 10.1063/1.464913.

[25] S. Grimme, J. Antony, S. Ehrlich, H. Krieg, A consistent and accurate *ab initio* parametrization of density functional dispersion correction (DFT-D) for the 94 elements H-Pu, J. Chem. Phys. 132 (2010) 154104. 10.1063/1.3382344.

[26] A.V. Marenich, C.J. Cramer, D.G. Truhlar, Universal Solvation Model Based on Solute Electron Density and on a Continuum Model of the Solvent Defined by the Bulk Dielectric Constant and Atomic Surface Tensions, J. Phys. Chem. B 113 (2009) 6378–6396. 10.1021/jp810292n.

[27] S. Dapprich, I. Komáromi, K.S. Byun, K. Morokuma, M.J. Frisch, A new ONIOM implementation in Gaussian98. Part I. The calculation of energies, gradients, vibrational frequencies and electric field derivatives, J. Mol. Struct. THEOCHEM 461–462 (1999) 1–21. 10.1016/S0166-1280(98)00475-8.

[28] T. Lu, A comprehensive electron wavefunction analysis toolbox for chemists, Multiwfn, J. Chem. Phys. 161 (2024) 082503. 10.1063/5.0216272.

[29] W. Humphrey, A. Dalke, K. Schulten, VMD: Visual molecular dynamics, J. Mol. Graph. 14 (1996) 33–38. 10.1016/0263-7855(96)00018-5.

[30] A. Canals, M. Purciolas, J. Aymamí, M. Coll, The anticancer agent ellipticine unwinds DNA by intercalative binding in an orientation parallel to base pairs, Acta Crystallogr. D Biol. Crystallogr. 61 (2005) 1009–1012. 10.1107/S0907444905015404.

[31] O. Trott, A.J. Olson, AutoDock Vina: Improving the speed and accuracy of docking with a new scoring function, efficient optimization, and multithreading, J. Comput. Chem. 31 (2010) 455–461. 10.1002/jcc.21334.

[32] Schrödinger, LLC, The PyMOL Molecular Graphics System, Version 1.8, (2015).

[33] M. Abraham, A. Alekseenko, B. Andrews, P. Bauer, C. Bergh, H. Bird, E. Briand, A. Brown, Y. Chen, M. Doijade, G. Fiorin, S. Fleischmann, S. Gorelov, G. Gouaillardet, A. Gray, F. Jalalypour, P. Johansson, C. Kutzner, G. Lazarski, J. Lemkul, M. Lundborg, J. Maia, P.T. Merz, V. Miletić, D. Morozov, L. Müllender, S. Páll, A. Pasquadibisceglie, M. Pellegrino, N. Piasentin, D. Rapetti, M.U. Sadiq, H. Santuz, M. Shirts, T. Shugaeva, A. Shvetsov, B. Soproni, P. Turner, A. Villa, Y. Zhang, B. Hess, E. Lindahl, GROMACS 2026.0 Source code, (2026). 10.5281/ZENODO.18299929.

[34] K. Lindorff-Larsen, S. Piana, K. Palmo, P. Maragakis, J.L. Klepeis, R.O. Dror, D.E. Shaw, Improved side-chain torsion potentials for the Amber ff99SB protein force field, Proteins Struct. Funct. Bioinforma. 78 (2010) 1950–1958. 10.1002/prot.22711.

[35] A.W. Sousa Da Silva, W.F. Vranken, ACPYPE - AnteChamber PYthon Parser interfacE, BMC Res. Notes 5 (2012) 367. 10.1186/1756-0500-5-367.

[36] G. Bussi, D. Donadio, M. Parrinello, Canonical sampling through velocity rescaling, J. Chem. Phys. 126 (2007) 014101. 10.1063/1.2408420.

[37] M. Parrinello, A. Rahman, Polymorphic transitions in single crystals: A new molecular dynamics method, J. Appl. Phys. 52 (1981) 7182–7190. 10.1063/1.328693.

[38] U. Essmann, L. Perera, M.L. Berkowitz, T. Darden, H. Lee, L.G. Pedersen, A smooth particle mesh Ewald method, J. Chem. Phys. 103 (1995) 8577–8593. 10.1063/1.470117.

39. [39] B. Hess, H. Bekker, H.J.C. Berendsen, J.G.E.M. Fraaije, LINCS: A linear constraint solver for molecular simulations, J. Comput. Chem. 18 (1997) 1463–1472. 10.1002/(SICI)1096-987X(199709)18:12%3C1463::AID-JCC4%3E3.0.CO;2-H.

[40] H. Sik Kim, S. Hyun Byun, B. Mu Lee, Effects of Chemical Carcinogens and Physicochemical Factors on the UV Spectrophotometric Determination of DNA, J. Toxicol. Environ. Health A 68 (2005) 2081–2095. 10.1080/15287390500182503.

[41] R. Bera, B.K. Sahoo, K.S. Ghosh, S. Dasgupta, Studies on the interaction of isoxazolcurcumin with calf thymus DNA, Int. J. Biol. Macromol. 42 (2008) 14–21. 10.1016/j.ijbiomac.2007.08.010.

[42] Beer, Bestimmung der Absorption des rothen Lichts in farbigen Flüssigkeiten, Ann. Phys. 162 (1852) 78–88. 10.1002/andp.18521620505.

[43] Y. Ni, S. Du, S. Kokot, Interaction between quercetin–copper(II) complex and DNA with the use of the Neutral Red dye fluorophor probe, Anal. Chim. Acta 584 (2007) 19–27. 10.1016/j.aca.2006.11.006.

[44] S. Roy, R. Banerjee, M. Sarkar, Direct binding of Cu(II)-complexes of oxicam NSAIDs with DNA backbone, J. Inorg. Biochem. 100 (2006) 1320–1331. 10.1016/j.jinorgbio.2006.03.006.

[45] G. Zhang, J. Guo, N. Zhao, J. Wang, Study of interaction between kaempferol–Eu3+ complex and DNA with the use of the Neutral Red dye as a fluorescence probe, Sens. Actuators B Chem. 144 (2010) 239–246. 10.1016/j.snb.2009.10.060.

[46] J.R. Lakowicz, ed., Principles of Fluorescence Spectroscopy, Springer US, Boston, MA, 2006. 10.1007/978-0-387-46312-4.

[47] N. Wang, L. Ye, B.Q. Zhao, J.X. Yu, Spectroscopic studies on the interaction of efonidipine with bovine serum albumin, Braz. J. Med. Biol. Res. 41 (2008) 589–595. 10.1590/S0100-879X2008000700007.

[48] MD. Hays, D.K. Ryan, S. Pennell, A Modified Multisite Stern−Volmer Equation for the Determination of Conditional Stability Constants and Ligand Concentrations of Soil Fulvic Acid with Metal Ions, Anal. Chem. 76 (2004) 848–854. 10.1021/ac0344135.

[49] P. Boguta, Z. Sokołowska, Zinc Binding to Fulvic acids: Assessing the Impact of pH, Metal Concentrations and Chemical Properties of Fulvic Acids on the Mechanism and Stability of Formed Soluble Complexes, Molecules 25 (2020) 1297. 10.3390/molecules25061297.

[50] S. Goutelle, M. Maurin, F. Rougier, X. Barbaut, L. Bourguignon, M. Ducher, P. Maire, The Hill equation: a review of its capabilities in pharmacological modelling, Fundam. Clin. Pharmacol. 22 (2008) 633–648. 10.1111/j.1472-8206.2008.00633.x.

[51] M.M.L. Fiallo, H. Tayeb, A. Suarato, A. Garnier-Suillerot, Circular Dichroism Studies on Anthracycline Antitumor Compounds. Relationship between the Molecular Structure and the Spectroscopic Data, J. Pharm. Sci. 87 (1998) 967–975. 10.1021/js970436l.

[52] S. Jaiswal, S.B. Dutta, D. Nayak, S. Gupta, Effect of Doxorubicin on the Near-Infrared OpticalProperties of Indocyanine Green, ACS Omega 6 (2021) 34842–34849. 10.1021/acsomega.1c05500.

[53] C. Guerra, J. Burgos, L. Ayarde-Henríquez, E. Chamorro, Formulating Reduced Density Gradient Approaches for Noncovalent Interactions, J. Phys. Chem. A 128 (2024) 6158–6166. 10.1021/acs.jpca.4c01667.

[54] S.R. Gadre, C.H. Suresh, N. Mohan, Electrostatic Potential Topology for Probing Molecular Structure, Bonding and Reactivity, Molecules 26 (2021) 3289. 10.3390/molecules26113289.

[55] K.M. Tewey, G.L. Chen, E.M. Nelson, L.F. Liu, Intercalative antitumor drugs interfere with the breakage-reunion reaction of mammalian DNA topoisomerase II., J. Biol. Chem. 259 (1984) 9182–9187. 10.1016/S0021-9258(17)47282-6.

[56] A. Nedělníková, P. Stadlbauer, M. Otyepka, P. Kührová, M. Paloncýová, Atomistic Insights Into Interaction of Doxorubicin With DNA : From Duplex to Nucleosome, J. Comput. Chem. 46 (2025) e70035. 10.1002/jcc.70035.

[57] V. Malatesta, A. Gervasini, F. Morazzoni, Chelation of copper(II) ions by doxorubicin and 4′-epidoxorubicin: ESR evidence for a new complex at high anthracycline/copper molar ratios, Inorganica Chim. Acta 136 (1987) 81–85. 10.1016/S0020-1693(00)87099-1.

[58] G. Das, A. Nicastri, M.L. Coluccio, F. Gentile, P. Candeloro, G. Cojoc, C. Liberale, F. De Angelis, E. Di Fabrizio, FT-IR, Raman, RRS measurements and DFT calculation for doxorubicin, Microsc. Res. Tech. 73 (2010) 991–995. 10.1002/jemt.20849.

[59] A. Jabłońska-Trypuć, G. Świderski, R. Krętowski, W. Lewandowski, Newly Synthesized Doxorubicin Complexes with Selected Metals—Synthesis, Structure and Anti-Breast Cancer Activity, Molecules 22 (2017) 1106. 10.3390/molecules22071106.

[60] H. Zhang, X. Chen, S. Qiao, H. Meng, H. Long, H. Zhong, Y. Liu, Y. Song, Y. Gao, Y. Liu, L. Mao, pH-Responsive Bovine Serum Albumin Nanoparticles Encapsulating Doxorubicin-Based Complexes Induce Cuproptosis in Lung Cancer Cells, Pharmaceutics 18 (2026) 526. 10.3390/pharmaceutics18050526.

[61] E. Todorović, S. Orzechowska, M. Milovanović, J. Pešić, M. Baranska, J.J. Lazarević, N. Lazarević, Temperature-induced spectral anomalies in doxorubicin—A Raman study, Spectrochim. Acta. A. Mol. Biomol. Spectrosc. 355 (2026) 127678. 10.1016/j.saa.2026.127678.

[62] S. Zhou, X. Feng, J. Bai, D. Sun, B. Yao, K. Wang, Synergistic effects and competitive relationships between DOC and DOX as acting on DNA molecules: Studied with confocal Raman spectroscopy and molecular docking technology, Heliyon 10 (2024) e30233. 10.1016/j.heliyon.2024.e30233.

[63] S. Nafisi, A.A. Saboury, N. Keramat, J.-F. Neault, H.-A. Tajmir-Riahi, Stability and structural features of DNA intercalation with ethidium bromide, acridine orange and methylene blue, J. Mol. Struct. 827 (2007) 35–43. 10.1016/j.molstruc.2006.05.004.

[64] M. Airoldi, G. Barone, G. Gennaro, A.M. Giuliani, M. Giustini, Interaction of Doxorubicin with Polynucleotides. A Spectroscopic Study, Biochemistry 53 (2014) 2197–2207. 10.1021/bi401687v.

[65] H. Xie, J. Jiang, X. Chu, H. Cui, H. Wu, G. Shen, R. Yu, Competitive interaction of the antitumor drug daunorubicin and the fluorescence probe ethidium bromide with DNA as studied by resolving trilinear fluorescence data: the use of PARAFAC and its modification, Anal. Bioanal. Chem. 373 (2002) 159–162. 10.1007/s00216-002-1306-y.

[66] H.-P. Xie, X. Chu, J.-H. Jiang, H. Cui, G.-L. Shen, R.-Q. Yu, Competitive interactions of adriamycin and ethidium bromide with DNA as studied by full rank parallel factor analysis of fluorescence three-way array data, Spectrochim. Acta. A. Mol. Biomol. Spectrosc. 59 (2003) 743–749. 10.1016/S1386-1425(02)00224-X.

[67] C. Ozluer, H.E.S. Kara, In vitro DNA binding studies of anticancer drug idarubicin using spectroscopic techniques, J. Photochem. Photobiol. B 138 (2014) 36–42. 10.1016/j.jphotobiol.2014.05.015.

[68] S.R. Madku, B.K. Sahoo, K. Lavanya, R.S. Reddy, A.T.S. Bodapati, DNA binding studies of antifungal drug posaconazole using spectroscopic and molecular docking methods, Int. J. Biol. Macromol. 225 (2023) 745–756. 10.1016/j.ijbiomac.2022.11.137.

